# Anti-resonance in developmental signaling regulates cell fate decisions

**DOI:** 10.1101/2025.02.04.636331

**Authors:** Samuel J. Rosen, Olivier Witteveen, Naomi Baxter, Ryan S. Lach, Erik Hopkins, Marianne Bauer, Maxwell Z. Wilson

## Abstract

Cells process dynamic signaling inputs to regulate fate decisions during development. While oscillations or waves in key developmental pathways, such as Wnt, have been widely observed, the principles governing how cells decode these signals remain unclear. By leveraging optogenetic control of the Wnt signaling pathway in both HEK293T cells and H9 human embryonic stem cells, we systematically map the relationship between signal frequency and downstream pathway activation. We find that cells exhibit a minimal response to Wnt at certain frequencies, a behavior we term anti-resonance. We developed both detailed biochemical and simplified hidden variable models that explain how anti-resonance emerges from the interplay between fast and slow pathway dynamics. Remarkably, we find that frequency directly influences cell fate decisions involved in human gastrulation; signals delivered at anti-resonant frequencies result in dramatically reduced mesoderm differentiation. Our work reveals a previously unknown mechanism of how cells decode dynamic signals and how anti-resonance may filter against spurious activation. These findings establish new insights into how cells decode dynamic signals with implications for tissue engineering, regenerative medicine, and cancer biology.

**Significance Statement:** Wnt signaling is responsible for driving stem cell differentiation. Our study explores a wide range of temporal patterns of Wnt activation using state-of-the-art optogenetics, advanced imaging and modeling. We identify anti-resonant frequencies that suppress mesoderm differentiation. We confirm this anti-resonant suppression for two human cell lines, HEK and embryonic stem cells, and expect that it also occurs in other pathways, as it arises from the interplay of different timescales along the pathway. Our work opens new avenues for systematically exploring signal spaces that natural systems can robustly respond to, and has broader implications for tissue engineering, regenerative medicine, and cancer biology.

## Introduction

Cells and tissues do not merely respond to static cues but process dynamic signals that encode crucial information about cell fate decisions. These signaling dynamics have gained significant attention due to the widespread use of live-cell fluorescent reporters that enable their visualization throughout a variety of developmental and cell fate transitions (1–3). For example, oscillations in conserved signaling pathways determine cell fate decisions in organisms ranging from yeast to human (4–6) and have been shown to selectively up-regulate specific genes (7, 8). In embryonic development it has become increasingly clear that dynamic signals encode information through waves (9–12), which are experienced by individual cells as periodic pulses or oscillations (13–16). Thus, understanding how cells decode dynamic signals is fundamental to unlocking insights into tissue development and regeneration.

In engineering, the concepts of resonance and anti-resonance illustrate how certain dynamic signals can be amplified or suppressed based on input frequency (17). When a system receives input at its resonant frequency, the response is amplified; in contrast, at an anti-resonant frequency, the response is diminished. While these principles have proven useful in engineering, their application to biological information processing remains largely unexplored. Recent advances in synthetic biology and cellular engineering have enabled the manipulation of signaling pathways with unprecedented precision, providing the opportunity to apply such engineering frameworks to cell fate decisions. Notably, optogenetic tools allow for reversible, rapid, and spatially confined activation of signaling pathways, creating a platform to explore the temporal dimension of signaling in high resolution (18). By systematically investigating the cellular responses to dynamical inputs we can uncover the ‘design principles’ underlying cell signaling and control, and apply these insights to optimize tissue engineering, regenerative therapies, and gain deeper insights into development.

A particular pathway that has both displayed a variety of dynamical signaling patterns (19) (pulses (20), oscillations (21, 22) and wave patterns (9, 23)) and that is involved in almost every developmental and regenerative cell fate decision is the Wnt signaling pathway. In addition to Wnt signals directing proliferation and differentiation of various adult stem cell niches, Wnt is canonically known for its role in specifying the primitive streak and the mesoderm germ layer during gastrulation of all higher vertebrates. Indeed, the Wnt pathway has a unique topology that suggests non-trivial, non-monotonic responses to dynamic inputs. Negative feedback has been described to act at every level of this pathway. At the receptor level, FZD/LRP6 receptors are internalized and degraded upon Wnt binding on the timescale of hours (24). In the cytoplasm, changes in inclusion of β-catenin (β-cat), the Wnt transcriptional effector, into Wnt processing biomolecular condensates (called the destruction complex/DC) occur on the timescale of 10s of minutes (25, 26). Finally, DC scaffold proteins, such as Axin, are transcribed and feed back onto DC activity on the timescale of hours (26). These layered feedback mechanisms suggest that the Wnt pathway can process a rich tapestry of temporal signals, making it a key candidate for investigating how cells interpret dynamic signals to drive precise developmental outcomes.

The Wnt pathway has been modeled as a system of ordinary differential equations (ODEs) (27, 28). Yet, these models involve many parameters (>20) and still fail to mechanistically capture the cell biology which plays a critical role in Wnt signal transduction. For example, Wnt signaling kinases and scaffold proteins form a phase-separated biomolecular condensate with a complex nucleation landscape which is not captured by simple ODEs that assume the components to be well-mixed (28). In contrast, abstracted models with a restricted number of “effective” or “hidden” variables are less reliant on the exact molecular wiring and instead focus on capturing the essential behaviors that emerge from underlying, often unobserved, interactions. This flexibility enables these abstracted models to generalize across different signaling pathways that more faithfully represent the input-output experimental framework enabled by the combination of optogenetic tools and live-cell reporters. This generality is particularly valuable in describing developmental signaling pathways, where their inherent versatility and complexity defy rigid, topologically fixed models.

Here we combine optogenetic control of the Wnt pathway with mechanistic and abstracted modeling to understand how the set of possible input dynamics are processed into cell fate decisions. We engineered a clonal cell line that contained both reversible, optogenetic control of the Wnt pathway as well as live-cell reporters that enabled precise, quantitative control and measurements of the Wnt signaling. By varying Wnt signal durations and monitoring responses, we explored Wnt target activity for simple optogenetic input patterns and developed an ODE model to fit their dynamics. The model predicts anti-resonant frequencies at which the response is suppressed. Experimental validation in HEK cells confirmed these predictions. We then built a minimal description of anti-resonance in the Wnt pathway, that abstracts the biochemical interactions into a single effective or hidden variable; we use the phrase “hidden variable” in analogy with hidden layers in neural networks. This model allows us to tune the properties of the anti-resonance using only two parameters. To demonstrate the generality and developmental relevance of our model, we engineered opto-Wnt into H9 human embryonic stem cells (hESCs). The mesoderm cell fate decision exhibited a stark dependence on stimulation dynamics, underscoring the developmental importance of anti-resonant frequencies. Overall, we present the first systematic optogenetic screen on developmental signaling dynamics in a mammalian stem cell line and pave the way to harnessing signaling dynamics for regenerative medicine and tissue engineering.

## Results

### A human cell line for all optical visualization and control of Wnt signaling dynamics

To enable simultaneous, precise control and real-time visualization of Wnt signaling, we engineered a clonal HEK293T cell line with optogenetic control over Wnt pathway dynamics by fusing Cry2 (29) to the LRP6 co-receptor (30, 31) (from here on referred to as the opto-Wnt tool). To track pathway outputs at multiple levels, we used a CRISPR knock-in strategy to endogenously tag β-catenin (β-cat) with a custom tdmRuby2 fluorescent protein, enabling real-time observation of transcription factor accumulation and degradation. Additionally, to monitor Wnt target gene transcription, we integrated an 8X-TOPFlash-tdIRFP (32) reporter through lentiviral transduction. This engineered cell line, designated as the Wnt I/O (Input/Output), is the first clonal cell line to offer both temporal control and live visualization of Wnt signaling, providing a platform for studying dynamic decoding in this pathway (Fig. 1A). We confirmed that both reporters respond to 450nm illumination through imaging and FACS (Fig 1B, Supp Fig. 1A).

**Figure 1.**
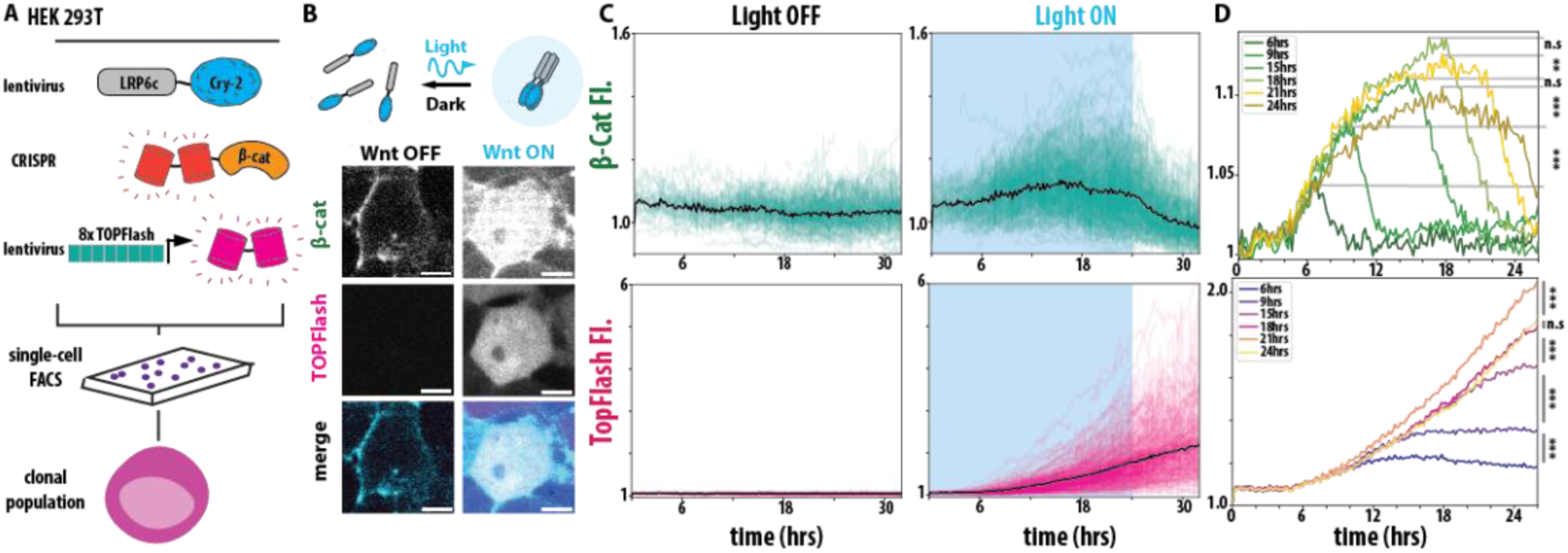
A human cell line for all optical visualization and control of Wnt signaling dynamics. A) Schematic of HEK293T Wnt I/O cells containing lentiviral optogenetic LRP6c-Cry2Clust, CRISPR tdmRuby3-β-cat, lentiviral 8X-TOPFlash-tdIRFP, clonally FACS-sorted. (B) Live cell imaging of HEK293T Wnt I/O cells exposed to no light and 24hrs of 405nm light illumination delivered every 2 minutes. Images are shown using the same lookup table. (C) Single-cell mean fluorescent intensity (MFI) traces (N = 321-567 cells, 4 biological replicates per condition) of tdmRuby3-β-cat and (TopFlash) tdiRFP measurements from live HEK293T Wnt I/O cells tracked during exposure to activating blue light (right) or no light (left) controls. Blue background indicates light on, white indicates light off. Black line represents population mean. (D) Population means of live, single-cell β-catenin (top) and TopFlash (bottom) MFI traces from indicated conditions (N = 321-595 cells, 4 biological replicates per condition, see Methods for significance values).

Next, given the observed timescales of Wnt-induced cell fate decisions, we performed a baseline ON-OFF experiment to understand both the dynamics and heterogeneity of the Wnt I/O line. We tracked single-cell β-catenin and TopFlash dynamics in over 300 single cells during 24 hrs of illumination to visualize pathway activation, followed by 8hrs of dark to visualize pathway de-activation (Fig. 1C, Supp. Mov 1). In comparison to the dark control, we noticed robust activation of the Wnt pathway on the population level, with variation in the activity patterns of individual cells. Notably, TopFlash reporter activity did not decay in the dark, while β-catenin showed relatively fast degradation in the dark. Wide variation in Wnt response among cell populations has been widely reported in both optogenetic and non-optogenetic cell lines (Supp Fig. 1B, C, 3, 26, 33). We attribute the observed cell population variation in both β-catenin and TopFlash to individual cells being in different phases of the cell cycle, which has been shown to influence Wnt signaling response (34, 35). To quantify our videos, individual cells were segmented and tracked between each frame of our videos using a custom image analysis pipeline that involved the CellPose-Trackmate (36) deep learning framework and focused on the nuclear fluorescence of β-catenin and TopFlash signals (Supp. Fig 1D, for details on tracking and segmentation see Methods). To reduce background noise in nuclear fluorescence signals, we first measured background fluorescence in the β-catenin and TopFlash channels for each frame. For each cell, raw nuclear β-catenin and TopFlash intensities were subtracted by the corresponding background value in that frame. The resulting traces were then normalized to the mean of all traces at t = 0. As a control, we also normalized the mean fluorescence of β-catenin and TopFlash in the light on condition to the light off condition (Supp Fig. 1E, F).

In addition to long duration Wnt signals (>24hrs), a wide range of shorter pulses of Wnt activity has been observed throughout mammalian development (21, 22). To understand the impact of these shorter Wnt pulses we performed a Wnt duration scan, varying the activation time between 6 and 21 hours, followed by a variable rest time for a total 26-hour experiment. We found that each cell population could be distinguished by either its maximum β-catenin concentration or the final level of TopFlash with statistical significance and that our choice of normalization had no impact on our results. (Fig. 1D, Supp. Table 1, Supp. Fig. 1G, H). We also observe that the TopFlash signal is delayed from β-catenin which responds earlier. Overall, these experiments demonstrate that our Wnt I/O line faithfully tracks and controls Wnt dynamics, allowing further interrogation of the signal processing capabilities.

Our quantifications reveal that β-catenin continues to accumulate in the presence of optogenetic stimulation, reaching a maximum at the end of optogenetic stimulation and decreases in the dark. Our statistical analysis shows a significant difference in the ‘light on’ maximum fluorescent intensity for β-catenin between most conditions with 18 and 21hrs showing no significance (Fig 1D, Supp. Fig. 1I). This observation is consistent with β-catenin saturating to a maximum level during a prolonged Wnt activation, as described by previous experiments and models of the Wnt pathways (37). Looking at the TopFlash response, we see that longer light durations led to higher TopFlash levels at the end of the experiment and that post-stimulus TopFlash levels increase monotonically with light duration. The results of our statistical analysis revealed that also all TopFlash conditions are statistically significant compared to one another (Fig 1D, Supp. Table 1, Supp. Fig. 1J). Overall, our data shows that our Wnt I/O cell line photo-switches on in light conditions and off in dark and can be used to quantify how temporal dynamics of Wnt signaling affect downstream gene expression.

### A model of Wnt signaling dynamics predicts anti-resonance

We then continued to analyze the ability of the Wnt signaling pathway to process dynamical inputs by developing a quantitative model. Although the Wnt pathway has been previously modeled, we sought to build a model of the Wnt pathway that captures our experimental observations with fewer parameters. We reduced the total number of differential equations in a previously established model of Wnt signaling (27, 38, 39) by describing only interactions between the DC and β-catenin (represented as state variables *c*(*t*) and *b*(*t*), respectively) (Fig. 2A). The complex formed when β-catenin binds to the DC is represented by state variable *c*_*b*_(*t*). We represent optogenetic Wnt activation (denoted as *l*(*t*)) as directly increasing disassociation of the DC, omitting receptor-level dynamics of the pathway and instead modeling the optogenetic response through the activation and deactivation of Dvl (30) (represented as the state variable *d*_*a*_(*t*) when active and *d*_*i*_(*t*) when inactive). In total, we have nine {*k*_1−7_, *d*_0_, *c*_0_} parameters governing DC and β-catenin dynamics, with *k*_1−7_ denoting rates and *d*_0_and *c*_0_denoting the conserved concentrations of Dvl and DC respectively (Methods and Supp. Text). Since our experiments do not track all state variables directly, we use literature values (28, 40–42) to eliminate five parameters, and constrain the remaining four parameters of the model using the experimental data (Methods and Supp. Text). We model TopFlash transcription (represented as the state variable *g*(*t*)) as a sigmoidal function of β-catenin accumulation. We also use experimental data to constrain the four free parameters {*r*_max_, *τ*, *n*, *K*} governing TopFlash expression, whose expression activates non-linearly, with a Hill function with parameters *n* and *K*, based on β-catenin levels at time *t* − *τ*. We arrived at the following differential equations to describe Wnt signalling dynamics:

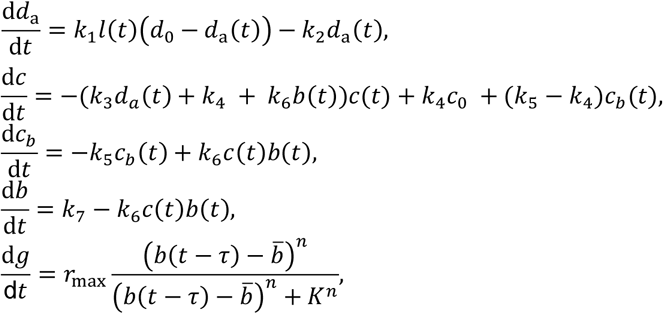

**Figure 2.**
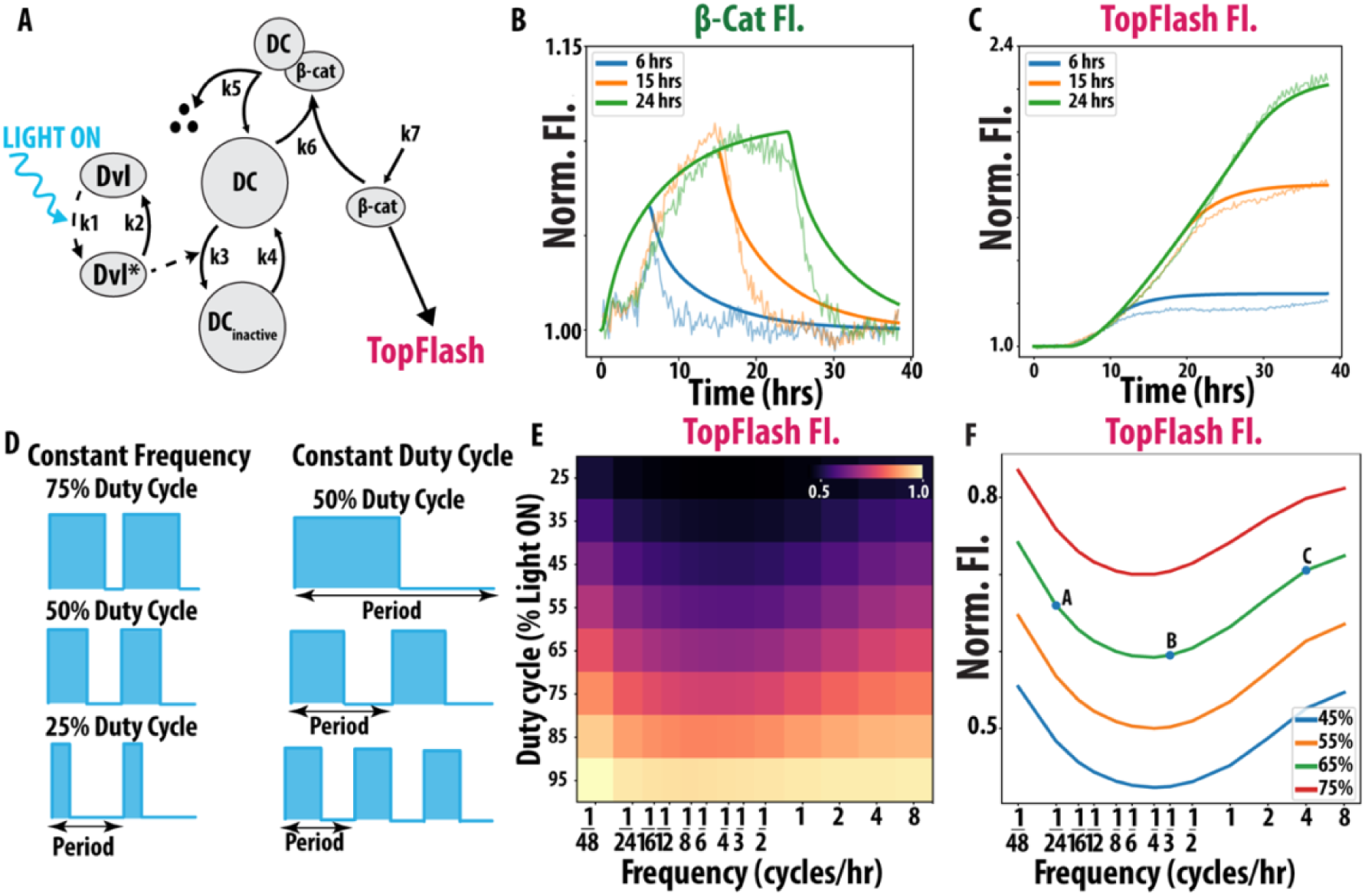
A model of Wnt signaling dynamics predicts anti-resonance. (A) Schematic of ordinary differential equation (ODE) model of Wnt signaling. Dotted lines represent light dependent parameters. For information on model variables and parameters, refer to Methods, ODE Model for Wnt Signaling. (B) ODE model predictions (solid) of β-cat mean fluorescent intensity (MFI) for 6, 15 and 24hrs compared against our unsmoothed experimental results (light) from Figure 1D. Post-26 hours, experimental data was corrected for over confluency effects in both β-cat and TopFlash. (C) ODE model predictions (solid) of TopFlash MFI compared against our unsmoothed experimental results (light) from Figure 1D. (D) Visualization of duty cycle and frequency. *Left:* Constant frequency with varying duty cycle. *Right:* Constant duty cycle with varying frequency. (E) ODE model generated heatmap of endpoint TopFlash MFI for various combinations of duty cycle and frequency conditions. F) Line graph of 45-75% duty cycles vs frequency with 1/24, 1/3 and 4 cycles/hr labeled as A, B and C.

where *b̅* denotes β-catenin at steady state when the light is off *l* = 0. We provide the complete list of parameters and their meaning in the Supplemental Text.

We compare our model’s β-catenin and TopFlash output for 6hr, 15hr and 24hrs of continuous light exposure to the same experimental conditions in Figure 1D. We observe that our model recapitulates our experiment results (Supp Fig. 1H, K) (solid v. lighter line) (Fig. 2B, C) with reasonably accuracy given cell-to-cell variability. This model therefore streamlines the complexity of Wnt signaling and still predicts β-catenin and TopFlash dynamics. While the mechanism proposed by the model has not been directly tested using experimental modification, such as specific inhibitors that perturb a single term in the model, our model agrees well with our experimental data. Assessing whether our model also reproduces the responses to these perturbations is an interesting direction for future work. We observe that our model matches TopFlash dynamics better than β-catenin dynamics. However, given the heterogeneity in expression profiles among the cell population and the use of literature values to constrain several parameters, our model still demonstrates a good agreement in both cases.

Using our model, we computationally explored the impact different complex temporal inputs have on Wnt signalling. First, we observe that TopFlash non-trivially accumulates in response to two Wnt signals of constant duration when the pause between them is varied (Supp. Text Section 1.2), reminiscent of non-trivial population dynamics outcomes under pulsed environmental variations (43). This prompted us to investigate how Wnt signaling is affected when keeping the total duration of the Wnt signal constant while varying the number of pulses per unit time.

We screened through a large region of Wnt input space that represents observed signalling dynamics during development (9, 10). We systematically varied the total integrated light exposure (duty cycle) and the number of pulses per unit time (frequency) (Fig. 2D). We simulated the β-catenin and TopFlash response of our Wnt model for 104 unique input combinations of duty cycle and frequency for 48hrs. We observe that β-catenin increases smoothly as frequency increases (Supp. Fig. 2B) for a given duty cycle. Notably, for a given duty cycle, the total level of TopFlash fluorescence is lowest at an intermediate frequency: we observe a minimum at ∼1/6 cycles/hr for all duty cycles (Fig 2E). Since TopFlash expression is reduced at these intermediate frequencies, we refer to this behavior as anti-resonance. Both the TopFlash heatmap as well as individual examples of constant duty cycle and varying frequency reveal a reduction in signal at intermediate frequencies consistent with anti-resonance (Fig. 2F). Individual temporal traces of TopFlash from experiment and simulation at low, high and anti-resonant frequencies (points A, B, C in Fig. 2F) also demonstrated the anti-resonant effect (Supp. Fig. 2C, D).

### The Wnt pathway of HEK cells displays anti-resonance

Next, we tested if the anti-resonant frequencies predicted by the model occur in the Wnt I/O cell-line. To simultaneously scan through a large range (=96) of unique duty cycle and frequency combinations, we utilized a high-throughput light stimulation device, the LITOS plate (44), that can deliver unique light patterns to all individual wells of a 96-well plate (Fig. 3A). Using this device, we performed the first ever optogenetic screen of Wnt pathway dynamics over a 48-hour period.

**Figure 3.**
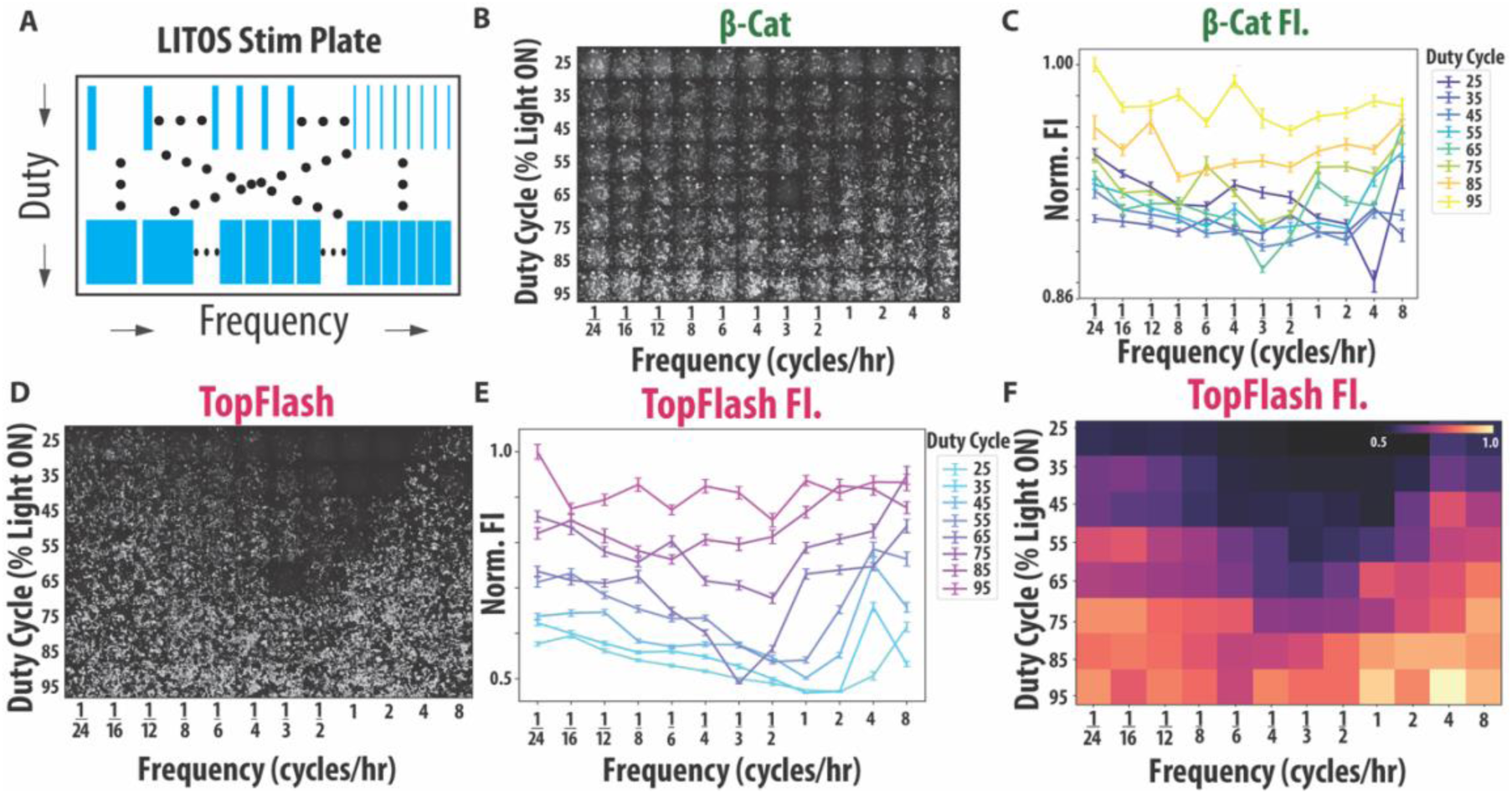
The Wnt pathway of HEK cells displays anti-resonance. (A) Schematic of our experimental method of the LITOS illumination device. (B) Qualitative images of end point β-catenin fluorescence post LITOS illumination with the heatmap of end point β-catenin MFI in the top right corner (N = 106-590 cells, 4 biological replicates per condition). (C) Error bar plot of end point β-catenin MFI post frequency and duty cycle screen. Error bars represent standard error of the mean (SEM). (D) Qualitative images of end point TopFlash fluorescence post LITOS illumination (N = 106-590 cells, 4 biological replicates per condition). (E) Error bar plot of end point TopFlash MFI post frequency and duty cycle screen. Error bars represent SEM. (F) Averaged heatmap of end point TopFlash MFI from two replicates of duty cycle and frequency experiment. Replicate heatmaps were normalized by the logarithm of the cell count at each well prior to averaging. Heatmap labels are displayed in categorical format, differentiating our experimental results heatmap from our computational heatmap.

We find that β-catenin is more uniformly expressed across frequencies than our model predicted, with some increase of β-catenin fluorescence for high frequency inputs (Supp. Fig. 2A, 3A, B). Based on our model, the β-catenin heatmap can be understood by the fast degradation times of β-cat (∼10 minutes) relative to the timepoint of the last pulse of this experiment, meaning that β-catenin fluorescence after 48h depends also on the pulse arrangement in a particular frequency/duty cycle combination (Fig. 3B, C, Supp. Fig. 3A, B). In addition to quantifying fluorescence, we confirmed that there were no statistical differences due to cell number or total integrated light exposure (Supp. Fig. 3C, D). Next, we look at TopFlash expression to verify the presence of the anti-resonance.

TopFlash displayed a clear anti-resonant effect with minimum activity between 1/3 and 1 cycles/hr, demonstrating that cells with the same duty cycle but different frequencies have varying pathway outputs (Fig. 3D, Supp. Fig. 3E). For example, the 55% duty cycle stimulation resulted in a clear decrease in TopFlash at the ∼1/2 cycles/hr frequency (Fig. 3E). We further verified robustness of our results by performing a replicate experiment where we rearranged the order of the duty cycle and frequency conditions, demonstrating that the anti-resonance is independent of the experimental arrangement (Supp. Fig. 3F). We quantified the TopFlash mean fluorescent intensity (MFI) of our replicate experiment and averaged them with our original experiment’s TopFlash MFI. Plotting the average of our two experiments as a heatmap, we observe a strong agreement between the results of our two experiments (Fig. 3F, Supp. Fig 3G, F). These experimental results confirm anti-resonance in Wnt signaling and further confirm the predictions by our computational model of the Wnt pathway.

### Hidden-variable approach reveals timescales of Wnt activation define shape of anti-resonance

We have shown that a biochemical model based on established models of the canonical Wnt pathway correctly predicts the anti-resonance observed in optogenetic Wnt activation. Next, we abstract our model further to establish a minimal model that explains this non-monotonic behavior. Rather than explicitly modelling the interactions of proteins upstream of β-catenin, we abstract the time-dependent response of the Wnt pathway into a single a “hidden variable” *a*(*t*) (Fig. 4A). This variable is activated optogenetically until saturation at a rate *k*_on_, and deactivated in absence of light at a rate *k*_off_. We use equivalent β-catenin dynamics to the ones earlier, with a degradation and a synthesis term, consistent with Goentoro et al (38). The degradation of β-catenin dynamics is inhibited by the hidden variable, such that the β-catenin accumulates during optogenetic stimulation as before. We describe TopFlash transcription using the same Hill-type activation as the biochemical model in Fig. 2 and note that the coefficient of the Hill-function has no significant effect on the presence of the anti-resonance.

**Figure 4.**
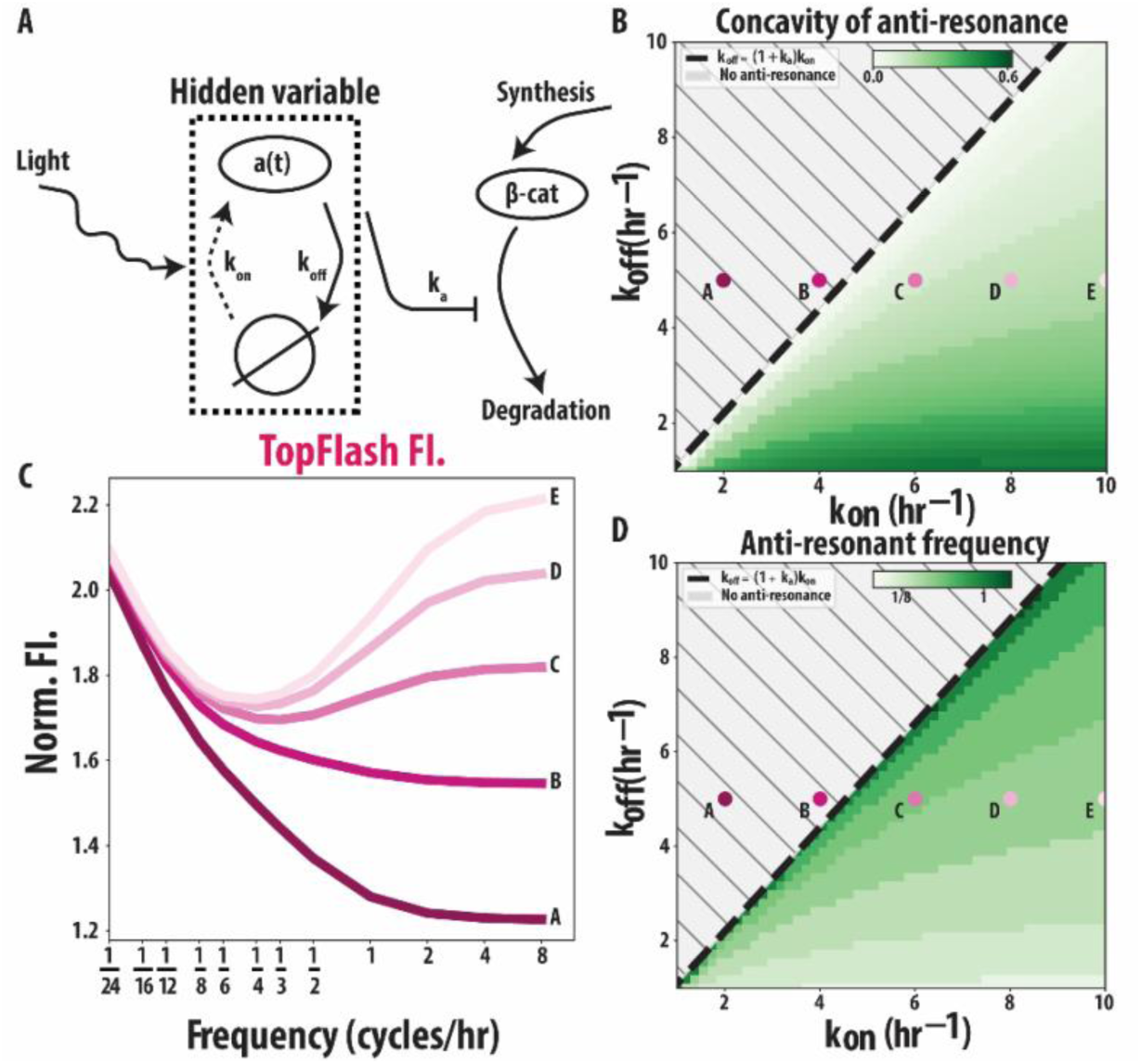
A hidden variable approach relates the anti-resonance to the timescales of Wnt activation and deactivation. (A) A hidden variable *a*(*t*) is activated upon optogenetic Wnt activation at a rate *k*_on_ and deactivates at rate *k*_off_ when the light is turned off. In turn, *a*(*t*) is coupled to first-order β-catenin dynamics *b*(*t*). (B-D) Systematic exploration of the parameter space shows that the rates *k*_on_ and *k*_off_ tune the concavity. (B) Concavity of anti-resonance is dependent on the combination of *k*_on_ and *k*_off_rates. (C) Shape of the anti-resonance for five different points (A-E) in parameter space. As we enter the region *k*_off_ < (1 + *k*_a_)*k*_on_ the anti-resonance appears, consistent with our analytical result (see Supplemental Text for details about equations and parameter values). (D) Anti-resonant frequency is dependent on the combination of *k*_on_ and *k*_off_ rates.

If the dynamics of the hidden variable occur at shorter timescales than the dynamics of β-catenin, we can reproduce the dynamics of β-catenin and TopFlash from Fig. 2 as well as the anti-resonance (see Supplemental Text). Our more abstract model allows us to see that the anti-resonance arises because of the interplay of the timescales involved. In the low frequency regime, the timescale of *a*(*t*) becomes irrelevant, and only the slower β-catenin dynamics are important. When the frequency is increased, the pulse duration is no longer long enough for β-catenin to saturate to a steady-state value. Especially when coupled with non-linear Hill-type activation, this leads to less TopFlash being produced. As the frequency increases further towards the high-frequency regime, the dynamics of *a*(*t*) become important. In this regime, we obtain increasing TopFlash expression with frequency as long as *k*_off_ < (1 + *k*_2_)*k*_on_ is satisfied. We confirm this bound analytically in Sec. 1.3 of the Supplemental Text. In Fig. 4B and D, we quantify the concavity of the anti-resonance and the corresponding anti-resonant frequency, respectively. As *k*_on_increases with *k*_off_ fixed, we observe that the minimum in TopFlash expression becomes sharper, while the location of the minimum shifts to the left. For five different points on the (*k*_on_, *k*_off_)-plane, we plot the final TopFlash level as a function of the frequency of the light cycles (Fig. 4C). We observe in Fig. 4B and 4C that the anti-resonant frequency indeed only appears once we enter the region *k*_off_ < (1 + *k*_2_)*k*_on_.

The explanation of anti-resonant behavior through two timescales in the hidden component(s) of the pathway suggest that the anti-resonance is a generic feature of the canonical Wnt pathway: it arises from differences in the timescale of activation and deactivation of the signaling cascade upstream of β-catenin. Mechanistically, the anti-resonant behavior is possible when the timescale of activation of the hidden variable is faster than its deactivation. In our full biochemical model, this can correspond, for example, to the phosphorylation timescale of Dvl protein at the receptor, which is thought to be fast for HEK293T cells (45).

Our minimal model suggests that the anti-resonant suppression of regulatory responses at pulses of intermediate frequencies does not require fine-tuning to specific rates, but it does require these rates to satisfy a loose bound. Therefore, the generality of the model suggests that this behavior may also occur in other cells.

### Anti-resonant dynamics drive mesodermal stem cell differentiation in hESC H9s

To investigate whether Wnt pathway anti-resonance influences stem cell fate decisions in the human embryo, we explored its role in the differentiation of H9 human embryonic stem cells (hESCs). Indeed, Wnt signaling pulses, oscillations, and waves have all been observed in organoid models of human development (e.g. gastruloids, somatoids, embryoid bodies) (46–48). Accordingly, we chose to address two questions: 1) Does anti-resonance affect the Wnt signals of human embryonic stem cells and, if so, (2) does it have a significant impact on developmental cell fate decisions?

We began by constructing a clonal H9 human embryonic stem cell (hESC) line containing our optoWnt tool and CRISPR-tagged tdmRuby β-catenin (Fig 5A). First, we CRISPR-tagged the N-terminus of β-catenin in H9 hESCs with tdmRuby. We next used the piggyBac transposon system to introduce our opto-Wnt tool into these H9 cells which were then selected for integrands and generated a clonal line (see Methods for details of cell line construction). Finally, we verified the opto-response by illuminating Wnt I/O H9s for 24 hours using 405nm light and staining for Brachyury (BRA), a mesodermal cell fate marker (49). Optogenetic stimulation of our Wnt I/O H9 cell line resulted in efficient mesodermal differentiation (Fig 5B). From these observations, we conclude that our Wnt I/O H9 line responds to optogenetic activation and that this activation can control the cell fate decision of mesoderm differentiation.

**Figure 5.**
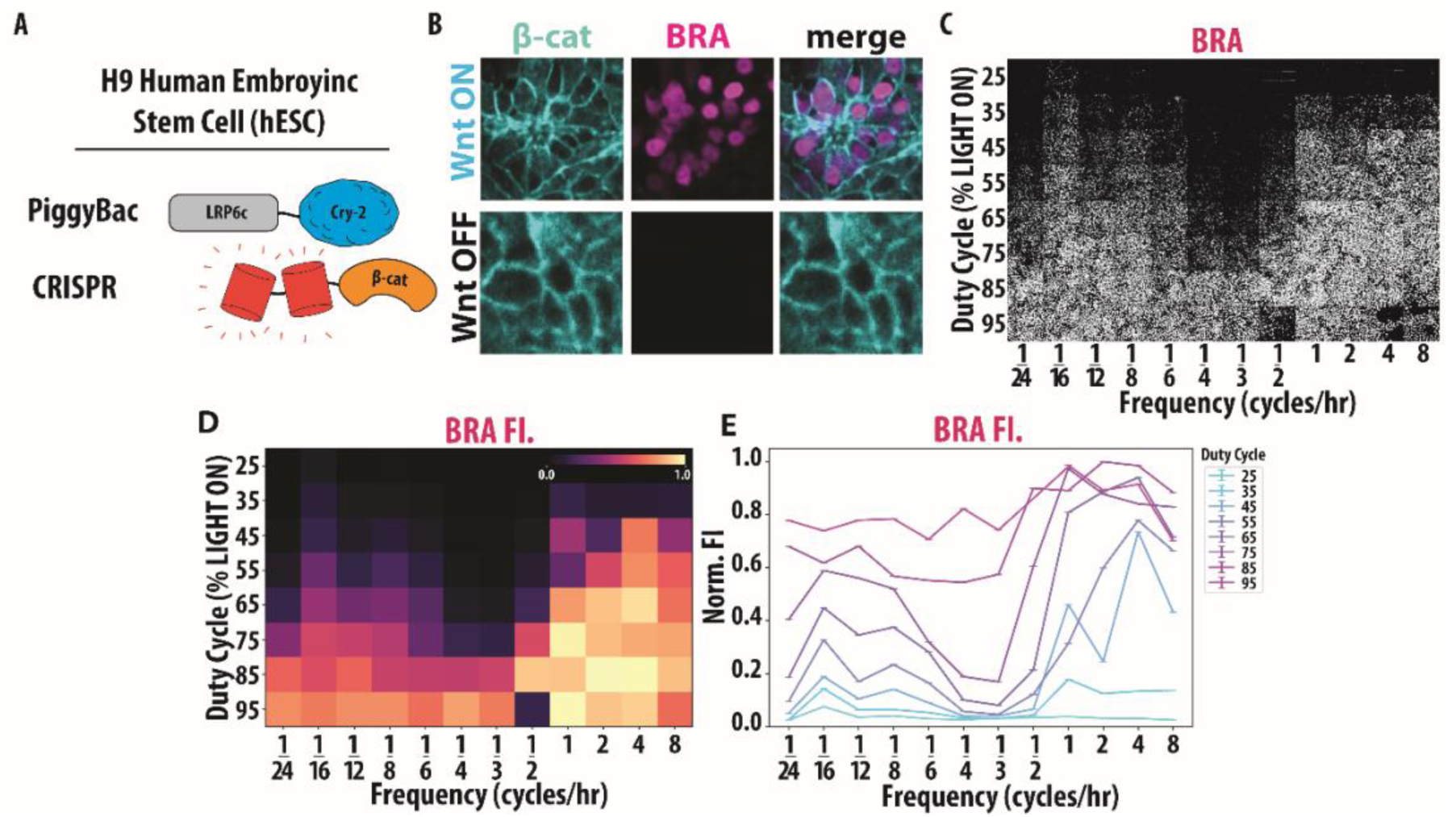
Anti-resonant dynamics drive mesodermal stem cell differentiation in hESC H9s. (A) Schematic of H9 Wnt I/O cells containing PiggyBac optogenetic LRP6c-Cry2Clust and CRISPR tdmRuby3-β-catenin. (B) Representative examples of tdmRuby3-β-cat and Brachyury (BRA) accumulating in response to 24hrs of blue light activation in H9 Wnt I/O cells, post puromycin selection. (C) Qualitative images of the end point Brachyury (BRA) fluorescence post LITOS illumination (N = 862-3176 cells, 6 biological replicates per condition). (D) Heatmap of end point BRA MFI for various duty cycle and frequency conditions. Heatmap labels are displayed in categorical format, differentiating our experimental results heatmap from our computational heatmap. (E) Error bar plot of end point BRA MFI post frequency and duty cycle experiment. Error bars represent standard error of the mean (SEM).

To determine whether anti-resonance affects mesoderm differentiation, we conducted a light stimulation screen targeting various duty cycles and frequencies, as described earlier (Fig. 3). We focused on a 24-hour differentiation window (50) as it is the literature reported timescale for Wnt driven mesoderm differentiation, and we found that using a 48-hour window caused all conditions to fully differentiate into mesodermal fates (Supp. Fig. 4A, B). We applied the same stimulation parameters previously tested in the HEK293T line and following the stimulation period, cells were fixed and stained (51) for BRA. We then fluorescently imaged our cells for both β-catenin and BRA (Fig 5C, Supp 4C, D, E). Quantification of β-catenin agrees well with our model and HEK293T results (Supp. Fig. 4F). Remarkably, quantification of BRA demonstrated a striking anti-resonant effect (Fig 5D). Namely, for cells with the same duty cycle (e.g. 55%), we observe that high and low frequencies induce total differentiation, while frequencies around 1/3 cycles per hour show markedly reduced BRA (Fig 5E). Together with our computational findings, these results demonstrate that anti-resonance is conserved across cell lines and that dynamic activation of the Wnt pathways influences stem cell differentiation in anti-resonant ways.

## Discussion

The concept that cells can interpret and respond to a diverse landscape of dynamic inputs has transformed our understanding of cellular signaling (7–16). Here, we explored this dynamic landscape by investigating the effects of periodic signals of varying frequency and duty cycle on the Wnt signaling pathway. Using a combination of computational, theoretical, and experimental approaches, we conducted the first optogenetic screen to probe the temporal dynamics of this essential signaling pathway in mammalian cells. We uncovered a previously unreported phenomenon in cellular signaling: for constant duty cycles, specific frequencies lead to significantly reduced levels of Wnt signaling. We called these anti-resonant frequencies, borrowing from engineering and electronics, where the term describes input frequencies that yield minimal system output. We confirmed the presence of the anti-resonance in both HEK293T cells and hESCs.

We speculate that the anti-resonance has biological meaning in that it suppresses a developmental response from oscillations with atypical timescales. For high-frequency oscillations, Wnt fluctuates rapidly, and it is likely not beneficial for the cells to respond to nuanced changes in fluctuations. For low frequencies, where the Wnt signal corresponds to pulses of multiple hours in duration, cells similarly require a strong and robust response. For intermediate frequencies, one might have expected a smooth interpolation between the low and high frequency regimes, but this does not occur; instead, the response to these intermediate frequencies is suppressed.

Analogous band-stop filtering should arise in other developmental circuits that couple a fast ‘ON’ step to slower deactivation or negative feedback. In Hedgehog, for example, PKA/CK1/GSK3-mediated partial proteolysis of Gli with slower recovery of full-length Gli creates the same fast-activation/slow-reset motif our hidden-variable model predicts will yield anti-resonance, and Wnt–Hedgehog crosstalk through the shared kinase GSK3 suggests such frequency selectivity could occur in other developmental signaling pathways (52). Especially in the context of Wnt, where individual receptors are internalized and degraded inside the cells for every binding ligand, requiring a detailed response to periodic inputs with fast periods would be cellularly costly (53, 54). Different timescales for activation and deactivation likely also occur in different pathways, including the Erk (55–57), NF-kB (58), BMP (59), Notch (60) and Hippo/Yap pathways. For some of these pathways, input oscillations determine differentiation, often also in non-trivial ways depending on the period of the oscillations (56, 57, 60). This suggests a non-trivial filtering of the input signals could be important in these pathways as well.

Anti-resonant frequencies could therefore represent a biological mechanism for noise filtering and signal discrimination. By suppressing specific input frequencies, cells can reduce spurious activation of downstream pathways, thereby improving the fidelity of information transmission. Here, we identify the anti-resonant frequency first in a Wnt reporter, whose expression is reduced at this frequency. When verifying its effect on the expression of the biologically meaningful differentiation reporter Bra, we observe a much stronger suppression at anti-resonance. In the context of oncogenesis, where intermediate Wnt is a hallmark of many cancers (61, 62) (e.g. the “Goldilocks Theory” (63)) anti-resonance may act as a natural suppressor of intermediate signaling. This raises the intriguing possibility that the timescales and negative feedback inherent to Wnt signaling incorporate mechanisms to suppress even the most pernicious oncogenic signals. Indeed, it has been previously reported that certain frequencies of Wnt activation lead to human stem cell apoptosis (33).

Our observation is analogous to anti-resonant systems in an engineering context, where the amplitude of the response is suppressed at a particular anti-resonant frequency. We would like to highlight that at the moment, the observation of this phenomenon requires optogenetic screens, as the fast and sharp changes in signal concentrations are difficult to check using microfluidics. Furthermore, light delivery hardware allows for the testing of large sets of dynamical signals simultaneously, enabling the discovery of anti-resonant inputs.

Our results suggest that anti-resonance arises from the interplay of distinct timescales governing the synthesis, degradation, and interactions of components within the Wnt pathway. We found that a set of five ODEs, which simplify an existing model of the Wnt pathway, predicted the existence of an anti-resonant frequency, which we confirmed in experiments. To gain further insight, we coarse-grained the degrees of freedom that we do not observe in the experiments into a single hidden variable and showed that this abstracted model reliably captures the anti-resonant behavior. This reduction, inspired by observing sloppy parameters (64), preserves the model’s core predictive features, enhances its interpretability and facilitates the extraction of meaningful insights. Here, we coarse-grained our model by considering the biochemical network and identifying the necessity for a hidden variable as a minimal effective model. Further work on how to consistently coarse-grain biochemical networks (65) to an input-output network with a minimum number of hidden or latent variables would provide a framework orthogonal and complementary to detailed modelling of molecular interactions. Here, this reduction allowed us to identify an interplay of timescales in the signaling pathway that makes possible the suppression of response to specific inputs. This increases our understanding how such pathways “compute” and what network architectures make these computations possible or efficient. We have shown here that the optogenetic setup presents a toolbox to investigate such hitherto unexplored input spaces.

### Outlook

Our work establishes an anti-resonant suppression of gene regulatory outputs for oscillatory signals at a frequency range. We connect this suppression to a hidden layer in the signaling network, which introduces an additional timescale to the problem. This consideration of different timescales is abundant in signaling cascades and the door to applying our approach to other signaling cascades with well-known dynamical responses, such as Erk (55–57), NF-kB (58), BMP (59), and Notch (66), which also exhibit oscillatory behavior and waves which can determine differentiation. Future work will explore whether anti-resonance is a universal feature of signaling networks and leverage this knowledge to uncover new regulatory mechanisms that may only become apparent in the context of highly dynamic signaling inputs.

## Materials and Methods

### Cell Lines

Human 293T cells were cultured at 37°C and 5% CO2 in Dulbecco’s Modified Eagle Medium, high glucose GlutaMAX (Thermo Fisher Scientific, 10566016) medium supplemented with 10% fetal bovine serum (Atlas Biologicals, F-0500-D) and 1% penicillin-streptomycin. Experiments in human Embryonic Stem Cell (hESC) lines were performed using the H9 hESC cell line purchased from the William K. Bowes Center for Stem Cell Biology and Engineering at UCSB. Cells were grown in mTeSR™ Plus medium (Stem Cell Technologies) on Matrigel® (Corning) coated tissue culture dishes and tested for mycoplasma in 2-month intervals.

### Wnt I/O 293T Cell Line generation

Clonal 293Ts containing CRISPR tdmRuby3-β-cat and oLRP6_Puro were obtained from Dr. Ryan Lach (26). Cells were co-transfected with pPig_8X-TOPFlash-tdIRFP_Puro obtained from Dr. Ryan Lach and Super PiggyBac Transposase (System Biosciences cat#: PB210PA-1) using manufacturers recommendations and standard PEI-based transfection procedures. Cells were incubated for 24hrs before replacing with fresh media. Cells were then subject to 12hrs of continuous 405nm light activation on the LITOS and single-cell FACS sorted for tdmRuby3+, tdIRFP+ into a 96-well plate. Cells were monitored for growth over 14 days, only wells containing single colonies (arising from attachment of a single clone) were kept for subsequent processing. Prospective clonal populations were then imaged pre- and post 12hr light activation to screen for low baseline expression of β-catenin and TOPFlash and medium-high expression post activation.

### Wnt I/O H9 Cell Line generation

Clonal β-catenin reporter lines were generated through CRISPR/Cas9-mediated homology directed repair using analogous methods to HEK293T counterparts. Accutase digested single hESCs were seeded onto Matrigel coated 12 well plates and transfected with Lipofectamine™ Stem Transfection Reagent (Invitrogen, STEM00015) according to manufacturer recommendations. Once the cells grew to confluency, they were selected with 2µg/mL puromycin in mTeSR plus. Clonal populations were isolated through single cell sorting with the SH800 Sony Cell Sorter and expanded in mTeSR™ Plus medium. The resulting population was screened and periodically treated with puromycin to ensure they mimicked canonical Wnt signaling upon optogenetic stimulation.

### Optogenetic Stimulation

Spatial patterning of light during timelapse fluorescent imaging sessions was accomplished via purpose-built microscope-mounted LED-coupled digital micromirror devices (DMDs) triggered via Nikon NIS Elements software. Stimulation parameters (brightness levels, duration, pulse frequency) were optimized to minimize phototoxicity while maintaining continuous activation of Cry-2. For DMD-based stimulation on the microscope, the final settings for ‘Light ON’ were 25% LED power (λ = 455nm), 2s duration pulses every 30s. For experiments that did not require frequent confocal imaging, cells were stimulated via a benchtop LED array purpose-built for light delivery to cells in standard tissue culture plates (‘LITOS’) (44). The same light delivery parameters were used for LITOS-based stimulation as for microscope mounted DMDs. Light was patterned to cover the entire surface of intended wells of plates used, rather than a single microscope imaging field.

CHIR and Wnt3a Stimulation. HEK293T cells were treated using 10µM of CHIR99201 (Stem Cell Technologies, 72052) or 2.5nM of Wnt3a ligand (R&D Systems, 5036-WN) added into Dulbecco’s Modified Eagle Medium, high glucose GlutaMAX (Thermo Fisher Scientific, 10566016) medium supplemented with 10% fetal bovine serum (Atlas Biologicals, F-0500-D) and 1% penicillin-streptomycin. Cells were then imaged with DAPI in the same media 16hrs post addition of CHIR99201 and Wnt3a ligand.

### Antibodies and Immunofluorescence

Primary antibodies used to stain for Brachyury and Sox2 in H9s were α-Sox2 (Cell Signaling 3579, 1:500 dil.) and α-Brachury (RnD AF2085, 1:500). Secondary antibodies used were α-Rbt-Alexa-488 (Thermofisher A21206, 1:1000) and α-Gt-Alexa-647 (Thermofisher A21447, 1:1000). Tissue fixation and staining was carried out using standard protocols using cold methanol (51). Immunofluorescent samples were imaged using confocal microscopy (see below). Nuclear stains were carried out using NucBlue Live ReadyProbes (Hoescht 33342, R37605) according to manufacturer’s instructions.

### Imaging

All live and fixed cell imaging experiments were carried out using a Nikon W2 SoRa spinning-disk confocal microscope equipped with incubation chamber maintaining cells at 37°C and 5% CO2. Glass-bottom culture plates (Cellvis # P96-1.5H-N) were pre-treated with bovine fibronectin (Sigma #F1141) in the case of 293Ts or Matrigel in the case of H9s, and cells were allowed to adhere to the plate before subsequent treatment or imaging. HEK293T cells were imaged in Dulbecco’s Modified Eagle Medium, high glucose GlutaMAX (Thermo Fisher Scientific, 10566016) medium supplemented with 10% fetal bovine serum (Atlas Biologicals, F-0500-D) and 1% penicillin-streptomycin. H9s were imaged in mTeSR™ Plus medium.

### Image Analysis

All quantification of raw microscopy images was carried out using the same general workflow: background subtraction > classification > measurement > normalization > statistical comparison. When possible, subcellular segmentation of nuclear fluorescence was performed via context-trained deep learning-based Cellpose 2.0 algorithm derived from the ‘nuclei’ or ‘cyto2’ pretrained models pre-packed with the current Cellpose software distribution available here: (1) (36). Single-cell tracking and raw measurements were performed with the ‘LAP Tracker’ function in the TrackMate plugin for ImageJ available here: https://imagej.net/plugins/trackmate/ (36, 67). Tracks containing fewer than 50 contiguous frames (spurious or exited camera field of view) were omitted from subsequent analysis. Mean fluorescent intensity of regions of interest were measured and subsequently processed. Raw measurements were compiled, processed, and plotted via custom Matlab (ver. R2024b) and Python (ver. 3.9.13) scripts, available upon request.

### Data Normalization for Cell Trajectories

We begin by using the segmented and tracked cells produced by our Image Analysis method above. Each segmented and tracked cell contains information of its nuclear β-catenin and TopFlash at each timepoint during our timelapse experiment. In order to reduce background noise in nuclear fluorescence signals, we first measured background fluorescence in the β-catenin and TopFlash channels for each frame. For each cell, raw nuclear β-catenin and TopFlash intensities were subtracted by the corresponding background value in that frame. The resulting traces were then normalized to their values at t = 0 such that all traces being at a normalized fluorescent intensity of 1.

### Data Normalization for Duty Cycle and Frequency Experiments

We begin by using the segmented cells produced by our Image Analysis method. We do not calculate tracks here since duty cycle and frequency screen experiments only include end point image so there is no temporal data. Each segmented and tracked cell contains information of its nuclear β-catenin and TopFlash. We measured background fluorescence in the β-catenin and TopFlash channels for each condition in the duty cycle and frequency experiment. Each segmented cell was background-subtracted. We then calculated the mean β-catenin and TopFlash fluorescence for each condition. Finally, we normalized each condition to the maximum mean β-catenin or TopFlash fluorescence, making the range of normalized fluorescent intensities between 0-1. For Figure 3F, background subtraction was performed on both the original and replicate experiments. Subsequently, the TopFlash fluorescence for each well was scaled by a factor of log(N), where N represents the cell count in that well. After applying this scaling, we calculated the mean between the original and replicate datasets to generate a combined dataset. Finally, the combined dataset was normalized to its maximum value before being visualized as a heatmap.

### ODE Model for Wnt Signaling

A simple model describing β-catenin accumulation in response to Wnt activation using a single differential equation, derived by Goentoro et al. (38), did not capture the results here. Adapting a more detailed model, for example Lee et al. (27) involving over 20 parameters. Thus, after parameter considerations detailed in the supplemental text, we arrived at the following differential equations describing β-catenin and TopFlash dynamics. Variables and parameters are defined in the main text and in Tables 1.1 and 1.2 below. The dynamics of β-catenin for a given Wnt input is governed by:

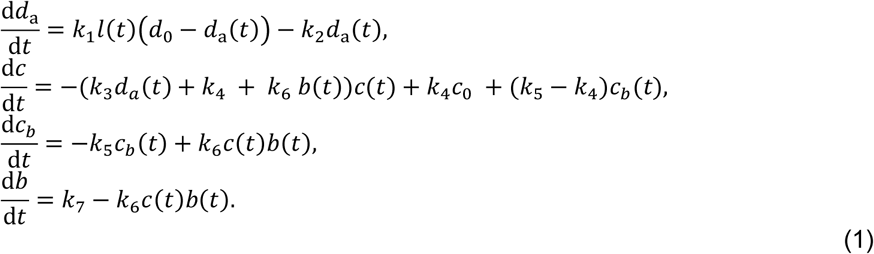

**Table 1.1.** Model variables.

| **variable** | **description** |
| --- | --- |
| $d_a$ | Active disheveled |
| $d_i$ | Inactive disheveled |
| $c$ | Free destruction complex |
| $c_b$ | β-catenin bound to destruction complex |
| $c_i$ | Inactive destruction complex |
| $b$ | Free β-catenin |
| $g$ | TopFlash expression |
| $l$ | Light status ($l \in \{0,1\}$) |

**Table 1.2.** Model parameters.

| **parameter** | **dimensions\*** | **description** |
| --- | --- | --- |
| $k_1$ | $\text{min}^{-1}$ | Rate of disheveled activation |
| $k_2$ | $\text{min}^{-1}$ | Rate of disheveled deactivation |
| $k_3$ | $\text{min}^{-1}$ | Inactivation rate of destruction complex by disheveled |
| $k_4$ | $\text{min}^{-1}$ | Activation rate of destruction complex |
| $k_5$ | $\text{min}^{-1}$ | Dissociation of phosphorylated β-catenin from the destruction complex |
| $k_6$ | $\text{min}^{-1}$ | Binding and phosphorylation of β-catenin with/by destruction complex |
| $k_7$ | $\text{min}^{-1}$ | β-catenin synthesis rate |
| $d_0$ | · | Conserved concentration of disheveled |
| $c_0$ | · | Conserved concentration of destruction complex |
| $n$ | · | Hill coefficient |
| $K$ | · | Dissociation constant |
| $r_{\max}$ | $\text{min}^{-1}$ | Maximum TopFlash transcription rate |
| $\tau$ | min | Time delay between β-catenin accumulation and onset of TopFlash transcription |

\* In our system of units, concentration is dimensionless.

In addition, we have the following conserved quantities:

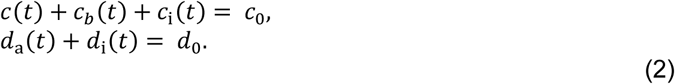

We model the dynamics of TopFlash using a simple Hill-type activation function:

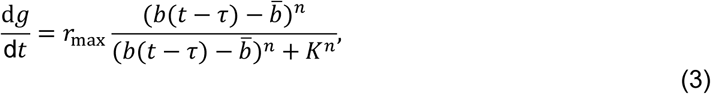

where we have added a time delay *τ* to represent the time delay between β-cat accumulation and TopFlash transcription.

### Abstract Model with Hidden Variable

We coarse-grain the model from Fig 2, described by Eqs. 1-3, to identify the core features of the model that give rise to the anti-resonance. The details of the signaling cascade upstream of β-catenin are coarse-grained into a single hidden variable *a*(*t*). We denote the presence of the stimulus at time *t* with *l*(*t*) ∈ {0,1}, with *l*(*t*) = 1 during “light on” and *l*(*t*) = 0 during “light off”. The variable *a*(*t*) is activated at rate *k*_on_ when *l*(*t*) = 1, and deactivated at rate *k*_off_ when *l*(*t*) = 0, i.e.:

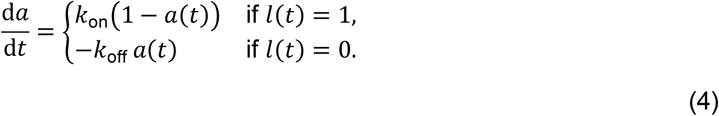

Note that we have normalized *a*(*t*) such that the steady state when *l*(*t*) = 1 is *a*(*t*) = 1. Next, we couple the hidden variable to the β-catenin dynamics *b*(*t*). We take it to be of the form derived by Goentero et al. (38):

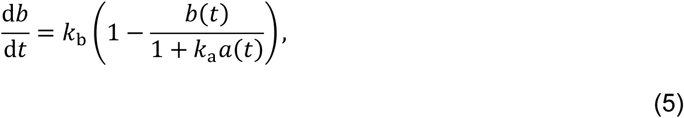

where *b*(*t*) has been rescaled so that the steady state when *l*(*t*) = 0 is *b*(*t*) = 1. Finally, we describe the TopFlash expression *g*(*t*) via the Hill-type activation function in Eq. 3 from the previous section (ODE model for Wnt Signaling).

The dynamical model described by Eqs. 3, 4, and 5 are summarized by the schematic in Fig. 4 in the main text. We confirm that this abstract model also fits the single-pulse data and refer to the Supplemental Text for more details. We find that parameters *k*_b_ and *k*_a_ dictate the coarse-grained β-catenin dynamics in Eq. 5, while the parameters *k*_on_ and *k*_off_govern the existence and shape of the anti-resonance. In the numerical simulations we observe that the anti-resonance occurs in a parameter regime *k*_off_ < (1 + *k*_a_)*k*_on_. We derive this bound in the Supplemental Text and sketch out the procedure here.

A periodic light input, like in our experiments, is determined by the duty cycle *δ* and frequency *f*, with period *T* = 1/*f*. For low frequencies *T*∼ 1/*k*_b_. we can ignore the comparatively fast changes in *a*(*t*) and replace it in Eq. 5 with *l*(*t*):

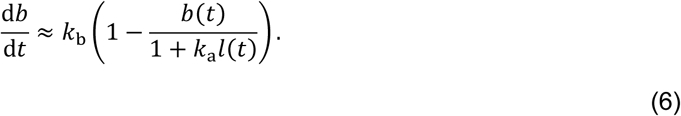

In this regime, the mean β-catenin level during the experiment strictly decreases as a function of frequency (see Supplemental Text). Since TopFlash expression approximately tracks mean β-catenin levels, it also decreases with frequency in this regime. We find that the exact functional form of TopFlash expression (Eq. 3), and the Hill parameter, influences the slope.

Next, we consider the high-frequency limit where *T* ≪ 1/*k*_b_. In this regime, the duration of each light pulse is short compared to the timescale in which significant changes occur in β-catenin levels. Hence, we can solve for small oscillations of beta-catenin Δ*b*(*t*) around a constant value *b̃*:

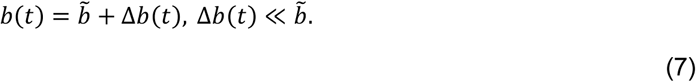

For a TopFlash reduction at intermediate frequencies, TopFlash should increase as the frequency increases towards the high-frequency limit. As the TopFlash activation function in Eq. 3 is strictly increasing in *b*(*t*), and Δ*b*(*t*) is small, it is sufficient to show that *b̃* increases as a function of frequency.

To first order in Δ*b*(*t*), the β-catenin dynamics in Eq. 5 can be written as:

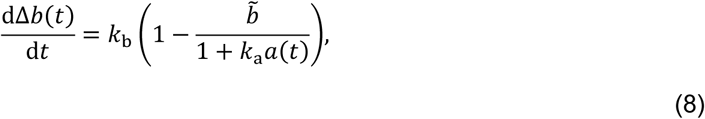

which we can solve for Δ*b*(*t*). Then, by imposing periodic boundary conditions, we can solve for the constant *b̃* (see Supplemental Text for details). For *b̃* to increase as the frequency increases towards the high-frequency limit, we require that:

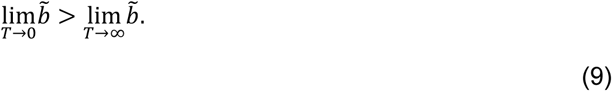

Substituting the above limits (derived in the Supplemental Text) yields the following condition:

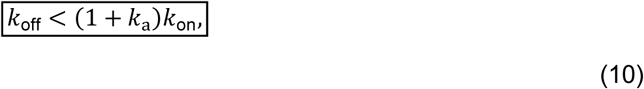

in agreement with results from the numerical simulations. All modeling code is available at is available at https://github.com/olivierwitteveen/wnt_antiresonance_model (68).

### Statistical Analysis

Statistical Analysis.
| p-value range | Symbol |
| --- | --- |
| p > 0.05 | n.s. |
| 0.01 < p ≤ 0.05 | *<br>— |
| 0.001 < p ≤ 0.01 | **<br>— |
| p ≤ 0.001 | ***<br>— |

## Acknowledgements

We acknowledge helpful discussion with E. F. Wieschaus, J. Rufo, A. Maynard, A. Bond and M. A. Morrissey.

M.B. acknowledges funding from the NWO Talent/VIDI program (NWO/VI.Vidi.223.169).

M.Z.W. acknowledges funding support from the NIH NICHD R01 HD108803-04.

## References

1. S. Regot, et al., High-Sensitivity Measurements of Multiple Kinase Activities in Live Single Cells. Cell 157, 1724–1734 (2014).

2. A. Yoney, et al., WNT signaling memory is required for ACTIVIN to function as a morphogen in human gastruloids. eLife 7, e38279 (2018).

3. S. Md. T. Rahman, J. M. Haugh, On the inference of ERK signaling dynamics from protein biosensor measurements. MBoC 34, ar60 (2023).

4. M. Matsuda, et al., Recapitulating the human segmentation clock with pluripotent stem cells. Nature 580, 124–129 (2020).

5. A. Mitchell, et al., Oscillatory stress stimulation uncovers an Achilles’ heel of the yeast MAPK signaling network. Science 350, 1379–1383 (2015).

6. P. Casani-Galdon, J. Garcia-Ojalvo, Signaling oscillations: Molecular mechanisms and functional roles. Current Opinion in Cell Biology 78 (2022).

7. M. Z. Wilson, et al., Tracing Information Flow from Erk to Target Gene Induction Reveals Mechanisms of Dynamic and Combinatorial Control. Molecular Cell 67, 757–769 (2017).

8. E. Batchelor, et al., Recurrent Initiation: A Mechanism for Triggering p53 Pulses in Response to DNA Damage. Mol Cell 30, 277–289 (2008).

9. I. Martyn, et al., A wave of WNT signaling balanced by secreted inhibitors controls primitive streak formation in micropattern colonies of human embryonic stem cells. Development 146 (2019).

10. S. Chhabra, et al., Dissecting the dynamics of signaling events in the BMP, WNT, and NODAL cascade during self-organized fate patterning in human gastruloids. PLoS Biol 17, e3000498 (2019).

11. P. Li, M. B. Elowitz, Communication codes in developmental signaling pathways Klein A. Development 146 (2019).

12. V. E. Deneke, S. Di Talia, Chemical waves in cell and developmental biology. Journal of Cell Biology 217, 1193–1204 (2018).

13. J. H. Levine, et al., Functional Roles of Pulsing in Genetic Circuits. Science 342, 1193–1200 (2013).

14. J. G. Albeck, et al., Frequency-Modulated Pulses of ERK Activity Transmit Quantitative Proliferation Signals. Molecular Cell 49, 249–261 (2013).

15. A. Aulehla, O. Pourquié, Oscillating signaling pathways during embryonic development. Current Opinion in Cell Biology 20, 632–637 (2008).

16. Y. el Azhar, et al., Unravelling differential Hes1 dynamics during axis elongation of mouse embryos through single-cell tracking. Development 151, dev202936 (2024).

17. V. Arora, Use of resonance and antiresonance frequencies for better matching of frequency response function. IJSTRUCTE 5, 13 (2014).

18. A. Möglich, K. Moffat, Engineered photoreceptors as novel optogenetic tools. Photochemical & Photobiological Sciences 9, 1286–1300 (2010).

19. J. van Zon, et al., Loss of Paneth cell contact starts a WNT differentiation timer in intestinal crypts. (2024).

20. T. Kroll JR, et al., Variability in β-catenin pulse dynamics in a stochastic cell fate decision in C. elegans. Developmental Biology 461, 110–123 (2020).

21. B. Mengel, et al., Modeling oscillatory control in NF-κB, p53 and Wnt signaling. Curr Opin Genet Dev 20, 656–664 (2010).

22. K. F. Sonnen, et al., Modulation of Phase Shift between Wnt and Notch Signaling Oscillations Controls Mesoderm Segmentation. Cell 172, 1079–1090.e12 (2018).

23. A. Hubaud, O. Pourquié, Signaling dynamics in vertebrate segmentation. Nature Reviews Molecular Cell Biology 15, 709–721 (2014).

24. M. V. Semënov, et al., DKK1 Antagonizes Wnt Signaling without Promotion of LRP6 Internalization and Degradation. Journal of Biological Chemistry 283, 21427–21432 (2008).

25. V. S. W. Li, et al., Wnt Signaling through Inhibition of β-Catenin Degradation in an Intact Axin1 Complex. Cell 149, 1245–1256 (2012).

26. R. S. Lach, et al., Nucleation of the destruction complex on the centrosome accelerates degradation of β-catenin and regulates Wnt signal transmission. Proc. Natl. Acad. Sci. U.S.A. 119, e2204688119 (2022).

27. E. Lee, et al., The Roles of APC and Axin Derived from Experimental and Theoretical Analysis of the Wnt Pathway. PLoS Biol 1, e10 (2003).

28. C. V. Giuraniuc, et al., A mathematical modelling portrait of Wnt signalling in early vertebrate embryogenesis. Journal of Theoretical Biology 551–552, 111239 (2022).

29. CRY2/CRY2. OptoBase. Available at: https://www.optobase.org/switches/Cryptochromes/CRY2-CRY2/

30. L. J. Bugaj, et al., Regulation of endogenous transmembrane receptors through optogenetic Cry2 clustering. Nat Commun 6, 6898 (2015).

31. C. Metcalfe, et al., Stability elements in the LRP6 cytoplasmic tail confer efficient signalling upon DIX-dependent polymerization. Journal of Cell Science 123, 1588–1599 (2010).

32. V. Korinek, et al., Constitutive Transcriptional Activation by a β-Catenin-Tcf Complex in APC^−/−^ Colon Carcinoma. Science 275, 1784–1787 (1997).

33. J. Massey, et al., Synergy with TGFβ ligands switches WNT pathway dynamics from transient to sustained during human pluripotent cell differentiation. Proceedings of the National Academy of Sciences 116, 4989–4998 (2019).

34. S. P. Acebron, et al., Mitotic Wnt Signaling Promotes Protein Stabilization and Regulates Cell Size. Molecular Cell 54, 663–674 (2014).

35. G. Davidson, et al., Cell Cycle Control of Wnt Receptor Activation. Developmental Cell 17, 788–799 (2009).

36. C. Stringer, et al., Cellpose: a generalist algorithm for cellular segmentation. Nature Methods 18, 100–106 (2021).

37. L. J. Bugaj, et al., Optogenetic protein clustering and signaling activation in mammalian cells. Nat Methods 10, 249–252 (2013).

38. L. Goentoro, M. W. Kirschner, Evidence that fold-change, and not absolute level, of beta-catenin dictates Wnt signaling. Mol Cell 36, 872–884 (2009).39.

39. S. M. De Man, et al., Quantitative live-cell imaging and computational modeling shed new light on endogenous WNT/CTNNB1 signaling dynamics. eLife 10, e66440 (2021).

40. K. Kang, et al., Dishevelled phase separation promotes Wnt signalosome assembly and destruction complex disassembly. Journal of Cell Biology 221, e202205069 (2022).

41. C. W. Tan, et al., Wnt Signalling Pathway Parameters for Mammalian Cells. PLoS ONE 7, e31882 (2012).

42. T. J. C. Harris, M. Peifer, Decisions, decisions: β-catenin chooses between adhesion and transcription. Trends in Cell Biology 15, 234–237 (2005).

43. M. Bauer, et al., Exploiting ecology in drug pulse sequences in favour of population reduction. PLoS Comput Biol 13, e1005747 (2017).

44. T. C. Höhener, LITOS: a versatile LED illumination tool for optogenetic stimulation. Sci Rep 12, 13139 (2022).

45. J. M. González-Sancho, et al, Wnt Proteins Induce Dishevelled Phosphorylation via an LRP5/6-Independent Mechanism, Irrespective of Their Ability To Stabilize β-Catenin. Molecular and Cellular Biology 24, 4757–4768 (2004).

46. E. Camacho-Aguilar, et al., Combinatorial interpretation of BMP and WNT controls the decision between primitive streak and extraembryonic fates. Cell Systems 15, 445–461.e4 (2024).

47. D.-L. Shi, Canonical and Non-Canonical Wnt Signaling Generates Molecular and Cellular Asymmetries to Establish Embryonic Axes. JDB 12, 20 (2024).

48. F. Aulicino, et al., Temporal Perturbation of the Wnt Signaling Pathway in the Control of Cell Reprogramming Is Modulated by TCF1. Stem Cell Reports 2, 707–720 (2014).

49. T. Faial, et al., Brachyury and SMAD signalling collaboratively orchestrate distinct mesoderm and endoderm gene regulatory networks in differentiating human embryonic stem cells. Development 142, 2121–2135 (2015).

50. M. Zhao, et al., Deciphering Role of Wnt Signalling in Cardiac Mesoderm and Cardiomyocyte Differentiation from Human iPSCs: Four-dimensional control of Wnt pathway for hiPSC-CMs differentiation. Sci Rep 9, 19389 (2019).

51. Immunocytochemistry protocol | Abcam.

52. M. Ding, X. Wang, Antagonism between Hedgehog and Wnt signaling pathways regulates tumorigenicity. Oncol Lett 14, 6327–6333 (2017).

53. X. Jiang, et al., Dishevelled Promotes Wnt Receptor Degradation through Recruitment of ZNRF3/RNF43 E3 Ubiquitin Ligases. Molecular Cell 58, 522–533 (2015).

54. H. Yamamoto, et al., Caveolin Is Necessary for Wnt-3a-Dependent Internalization of LRP6 and Accumulation of β-Catenin. Developmental Cell 11, 213–223 (2006).

55. K. M. Waters, et al., ERK Oscillation-Dependent Gene Expression Patterns and Deregulation by Stress Response. Chem. Res. Toxicol. 27, 1496–1503 (2014).

56. M. Yadav, et al., Homeorhetic regulation of cellular phenotype. [Preprint] (2025). Avaliable at: https://www.biorxiv.org/content/10.1101/2025.06.06.658216v1.

57. H. Ryu, et al., Frequency modulation of ERK activation dynamics rewires cell fate. Mol Syst Biol 11, 838 (2015).

58. D. E. Nelson, et al., Oscillations in NF-κB Signaling Control the Dynamics of Gene Expression. Science 306, 704–708 (2004).

59. S. Teague, et al., Time-integrated BMP signaling determines fate in a stem cell model for early human development. Nat Commun 15, 1471 (2024).

60. S. D. C. Weterings, et al., NOTCH-driven oscillations control cell fate decisions during intestinal homeostasis. [Preprint] (2024). Available at: https://www.biorxiv.org/content/10.1101/2024.08.26.609553v2

61. H. Zhao, et al., Wnt signaling in colorectal cancer: pathogenic role and therapeutic target. Molecular Cancer 21 (2022).

62. T. Zhan, et al., Wnt signaling in cancer. Oncogene 36, 1461–1473 (2017).

63. C. Albuquerque, et al., The “just-right” signaling model: APC somatic mutations are selected based on a specific level of activation of the beta-catenin signaling cascade. Human Molecular Genetics 11, 1549–1560 (2002).

64. R. N. Gutenkunst, et al., Universally Sloppy Parameter Sensitivities in Systems Biology Models. PLoS Comput Biol 3, e189 (2007).

65. M. Xia, et al., Structured Pruning Learns Compact and Accurate Models. [Preprint] (2022). Available at: http://arxiv.org/abs/2204.00408.

66. M. J. van Oostrom, et al., Coupling of cell proliferation to the segmentation clock ensures robust somite scaling. [Preprint] (2025). Available at: https://www.biorxiv.org/content/10.1101/2025.01.10.632257v2.

67. D. Ershov, et al., TrackMate 7: integrating state-of-the-art segmentation algorithms into tracking pipelines. Nat Methods 19, 829–832 (2022).

68. O. Witteveen, olivierwitteveen/wnt_antiresonance_model: v1.0.1 Anti-resonance in the Wnt pathway. (2025).

