## Supplemental Text for "Anti-resonance in developmental signaling regulates cell fate decisions"

**1. Model for β-catenin and TopFlash dynamics**

The results from Fig. 1D show the β-catenin (β-cat) and TopFlash dynamics in response to a single pulse of light. Figure 3 explores more complex patterns of light pulses, revealing a non-monotonic relationship between the dynamics of LRP6 activation and the level of TopFlash expression. Our goal is to formulate a minimal biochemical model that captures the β-cat and TopFlash dynamics observed in Fig. 1 and correctly predicts the anti-resonance observed in Fig. 3.

Quantitative measurements of parameters in the canonical Wnt pathway are not widely available[1–3]. The first computational model for Wnt signaling was developed for Xenopus egg extracts by Lee et al.[1] and has been widely used to quantify Wnt pathway dynamics in other systems. This model relies on parameters detailing protein–protein interactions, protein synthesis/degradation, and phosphorylation/dephosphorylation, as well as concentrations of key signaling proteins. However, these protein abundances can vary significantly between Xenopus extracts and mammalian cell lines[2], and hence it is difficult to apply this Xenopus model directly to our cells. Thus, we will develop a simple model in the following. We emphasize that our objective here is not to precisely calibrate biophysical parameters for the canonical Wnt pathway in HEK293T cells, but rather to show that the observed dynamics can emerge from known interactions within the pathway.

Modeling of β-cat dynamics is challenging as the protein is distributed between cytoplasmic and nuclear pools. Further, β-cat is an important part of the cell-to-cell adhesion complex[4]. The mechanism through which Wnt signaling affects subcellular distribution of β-cat is not fully understood. For the purposes of simplifying the model, we do not explicitly model β-cat localization.

We first attempt a model that captures β-cat dynamics on the most-coarse grained level using a single differential equation:

$$\frac{\mathbf{d}\boldsymbol{b}}{\mathbf{d}\boldsymbol{t}} = \boldsymbol{\alpha_1} - \frac{\boldsymbol{\alpha_2}\, \boldsymbol{b}(\boldsymbol{t})}{\mathbf{1} + \boldsymbol{\alpha_3} \boldsymbol{l}(\boldsymbol{t})}, \tag{1}$$

where $\{\boldsymbol{\alpha}_\mathbf{i}\}$ are rate constants, $\boldsymbol{b}(\boldsymbol{t})$ denotes the β-cat concentration, and $\boldsymbol{l}(\boldsymbol{t}) \in \{\mathbf{0}, \mathbf{1}\}$ is the light signal. The first and second term on the right-hand side capture β-cat synthesis and DC-dependent degradation, respectively. The latter is turned “off” by the Wnt signal. This physically intuitive equation was derived by Goentoro et al.[5] from the Xenopus model[1] using separation of timescales. We find that this simple model can capture β-cat dynamics of Fig. 1 reasonably well. However, we find that it does not capture well the anti-resonance in Fig. 3; even when coupled to non-linear TopFlash activation functions, we observed only weak effects for highly specific parameter combinations. Hence, we decided to extend the model.

We add a layer of complexity to the model by considering destruction complex (DC) dynamics. We coarse-grain the DC formation-cycle by treating it as an assembled object, where the light causes the DC to break down into an “inactivated” form. Binding of β-cat with the DC leads to the release of phosphorylated β-cat and recycling of the DC, and phosphorylated β-cat is subsequently degraded. Finally, we model TopFlash expression using a Hill-type function of the β-cat concentration. We find that this model phenomenologically captures the dynamics in Fig. 1

and Fig. 3. However, we noticed that the anti-resonance in Fig. 3 could be better described by including additional upstream protein dynamics. We will address this latter observation in Sec. 1.3. With this added step, stimulation by light causes activation of the disheveled (Dvl) protein instead of affecting the DC directly, as shown in Fig. 2A.

Thus, we arrive at the following differential equations describing β-cat and TopFlash dynamics. We define the variables and parameters in Tables 1.1 and 1.2 shown in Methods, but note that we typically use "$k$" to refer to rates and "$b$", "$c$", and "$d$" to refer to β-catenin, DC, and Dvl concentrations, respectively.

The dynamics of β-cat for a given Wnt input is governed by:

$$\begin{aligned}
\frac{\mathrm{d}d_{\mathrm{a}}}{\mathrm{d}t} &= k_1 l(t)(d_0 - d_{\mathrm{a}}(t)) - k_2 d_{\mathrm{a}}(t),\\
\frac{\mathrm{d}c}{\mathrm{d}t} &= -(k_3 d_a(t) + k_4 + k_6\, b(t))c(t) + k_4 c_0 + (k_5 - k_4)c_b(t),\\
\frac{\mathrm{d}c_b}{\mathrm{d}t} &= -k_5 c_b(t) + k_6 c(t) b(t),\\
\frac{\mathrm{d}b}{\mathrm{d}t} &= k_7 - k_6 c(t) b(t).
\end{aligned} \tag{2}$$

In addition, we have the following conserved quantities:

$$\begin{aligned}
c(t) + c_b(t) + c_{\mathrm{i}}(t) &= c_0,\\
d_{\mathrm{a}}(t) + d_{\mathrm{i}}(t) &= d_0.
\end{aligned} \tag{3}$$

Like in the main manuscript, $d_{\mathrm{i}}$ and $c_{\mathrm{i}}$ denote the inactive forms of Dvl and the DC, respectively, and $c_b$ denotes the β-catenin bound form of the DC. We model the dynamics of TopFlash using a simple Hill-type activation function:

$$\frac{\mathrm{d}g}{\mathrm{d}t} = r_{\max} \frac{(b(t-\tau) - \bar{b})^n}{(b(t-\tau) - \bar{b})^n + K^n}, \tag{4}$$

where we have added a time delay $\tau$ to represent the time delay between β-cat accumulation and TopFlash transcription.

Our model, while being significantly more compact compared to the model for Xenopus[1], correctly predicts non-trivial dynamics of TopFlash and β-cat accumulation in Figs. 1 and 3. It shares similarities with the model presented in de Man et al[3]. However, contrary to the latter, we do not consider shuttling of β-cat between the cytoplasm and nucleus, or explicitly model interactions of β-cat with transcriptional co-activators. In our experiments, we did not quantify the dynamics of the DC directly. Hence, we use a combination of experimental data and values taken from literature to constrain the parameters of our minimal model, as we outline in Section 1.1.

*1.1 Model analysis:*

We calculate steady state solutions and use these to fix parameter values, as discussed in the sections below.

*1.1.1 Steady state solutions:*

Setting all time derivatives in Eqs. (2) to zero, we can solve for the steady state of the model. Let us denote the steady state concentration with a horizontal bar. Setting $\boldsymbol{l} = \mathbf{0}$ or $\boldsymbol{l} = \mathbf{1}$ yields the steady-state solutions for light "off" and "on", respectively:

$$\begin{aligned} \overline{d_a} &= \frac{k_1 l}{k_1 l + k_2} d_0, \\ \bar{c} &= \left(c_0 - \frac{k_7}{k_5}\right) \frac{k_4}{\overline{d_a}\, k_3 + k_4}, \\ \overline{c_b} &= \frac{k_7}{k_5}, \\ \bar{b} &= \frac{k_7}{k_6} \frac{1}{\bar{c}}. \end{aligned} \tag{5}$$

*1.1.2 Parameter values*

Here, we use a combination of our experimental data and values obtained from literature to constrain the parameters of the model.

We work in dimensionless units of concentration where $\bar{b} = 1$ when the light is off, $l = 0$. This allows us to write:

$$c_0 = k_7 \left(\frac{1}{k_5} + \frac{1}{k_6}\right). \tag{6}$$

Further, we estimate from Fig. 1 that $\bar{b} \approx 1.1$ when $l = 1$. This implies:

$$k_3 \approx 0.10\, \frac{k_4}{d_0} \left(1 + \frac{k_2}{k_1}\right). \tag{7}$$

Next, we assume that the ratio $\bar{b}/\bar{c}$ when $l = 0$ is similar to de Man et al[3]. This yields:

$$k_6 \approx \frac{91.0\ \text{nM}}{82.4\ \text{nM}} k_7 \approx 1.1\, k_7. \tag{8}$$

We also assume that the ratio $\bar{b}/\bar{c_b}$ is similar to de Man et al.[3] This yields:

$$k_5 \approx \frac{91.0\text{ nM}}{62.5\text{ nM}} k_7 \approx 1.5\,k_7. \tag{9}$$

Finally, we assume the β-cat synthesis rate reported in Lee et al[1]. We convert to our dimensionless units of concentration by using $\bar{b} = 91$ nM from de Man et al[3]. We obtain:

$$k_7 \approx \frac{0.423\text{ nM min}^{-1}}{91\text{ nM}} \approx 4.6 \times 10^{-3}\text{ min}^{-1}. \tag{10}$$

We also choose to set $d_0 = 1$. For the remaining parameters $k_1$, $k_2$, and $k_4$, we run a numerical least-squares optimization using the data for single pulses. We find that a range of values of $k_1$, $k_2$ and $k_4$ capture the dynamics and use $k_1 = 0.17\ \mathbf{min}^{-1}$, $k_2 = 0.17\ \mathbf{min}^{-1}$, and $k_4 = 0.050\ \mathbf{min}^{-1}$ for the rest of the manuscript. For the TopFlash dynamics, we use $n = 2$, $r_{\max} = 0.11$ min$^{-1}$, $K = 1.0$, and $\tau = 4.0$ **hrs**.

Our model matches the data well for both TopFlash and β-catenin, especially given the single-cell variability. We observe that the match in beta-catenin could have been improved further if we had added an additional time delay in its transcription; since previous work did not incorporate this delay, we chose to not capture this delay here and instead add it for TopFlash expression.

*1.2 Behavior of the model for two pulses:*

Next, we explore the behavior of the model for two light pulses when we vary the pause duration. We expected a non-trivial response to pairs of pulses, as population dynamics model systems exposed to pulse sequences can show non-trivial responses; for example, when bacterial populations are exposed to antibiotics, two shorter antibiotic pulses can achieve the same response as a long one (see e.g. Bauer et al.[6]). We find that for short pauses $\lesssim 1$ hour, the final level of TopFlash expression does not change significantly (see Supp. Fig. 5). For longer pauses, TopFlash expression decreases when the pause duration is increased. When a very long pause is present, the final TopFlash expression is about ~30% lower compared to when there is no pause. The observed effect could be of importance, as it means that the organism does not need to sustain Wnt activation continuously to obtain the same differentiated output.

*1.3 Anti-resonance via a hidden variable:*

We observe an "anti-resonance" effect when the cells are subjected to periodic Wnt signals. That is, for the same amount of total integrated Wnt signal, intermediate frequencies result in the lowest amount of gene expression (see Fig. 3 and Fig. 5 in the main text). We found that a simple biochemical model based on existing models of the canonical Wnt pathway, with parameters values from literature, correctly predicts the anti-resonance (see Fig. 2 in the main text). By further coarse graining the model, we aim to identify the core features of the model that give rise to the anti-resonance. In what follows, we seek to place constraints on model parameters for the anti-resonance effect to occur.

The canonical Wnt pathway has a time-dependent response – it takes time before changes in the light status propagate through the signaling cascade and affect β-cat levels. Intermediate steps include, for example, activation of disheveled protein and inactivation of the destruction complex. Rather than modelling these steps explicitly, we capture the time-dependent response of the pathway via a "hidden variable" $\boldsymbol{a(t)}$. Let us denote the light status with $\boldsymbol{l(t)} \in \{\mathbf{0}, \mathbf{1}\}$, where $\boldsymbol{l(t)} = \mathbf{0}$ and $\boldsymbol{l(t)} = \mathbf{1}$ correspond to "light off" and "light on" respectively. Let $\boldsymbol{a(t)}$ be activated at rate $\boldsymbol{k}_{\text{on}}$ when $\boldsymbol{l(t)} = \mathbf{1}$, and deactivated at rate $\boldsymbol{k}_{\text{off}}$ when $\boldsymbol{l(t)} = \mathbf{0}$. The first-order dynamics of $\boldsymbol{a(t)}$ are described by:

$$\frac{\mathbf{d}\boldsymbol{a}}{\mathbf{d}\boldsymbol{t}} = \begin{cases} \boldsymbol{k}_{\text{on}}\big(\mathbf{1} - \boldsymbol{a(t)}\big) & \text{if } \boldsymbol{l(t)} = \mathbf{1}, \\ -\boldsymbol{k}_{\text{off}}\,\boldsymbol{a(t)} & \text{if } \boldsymbol{l(t)} = \mathbf{0}. \end{cases} \tag{11}$$

Note that we have normalized $\boldsymbol{a(t)}$ such that the steady state when $\boldsymbol{l(t)} = \mathbf{1}$ is $\boldsymbol{a(t)} = \mathbf{1}$. Next, we couple the hidden variable to the β-cat dynamics $\boldsymbol{b(t)}$. We take it to be of the form derived by Goentoro et al.[5]:

$$\frac{\mathbf{d}\boldsymbol{b}}{\mathbf{d}\boldsymbol{t}} = \boldsymbol{k}_{\mathbf{b}}\left(\mathbf{1} - \frac{\boldsymbol{b(t)}}{\mathbf{1} + \boldsymbol{k}_{\mathbf{a}}\boldsymbol{a(t)}}\right), \tag{12}$$

where $\boldsymbol{b(t)}$ has been rescaled so that the steady state when $\boldsymbol{l(t)} = \mathbf{0}$ is $\boldsymbol{b(t)} = \mathbf{1}$. Finally, we describe the TopFlash expression $\boldsymbol{g(t)}$ via a Hill-type activation function like before:

$$\frac{\mathbf{d}\boldsymbol{g}}{\mathbf{d}\boldsymbol{t}} = \boldsymbol{r}_{\text{max}}\frac{(\boldsymbol{b(t-\tau)} - \mathbf{1})^2}{(\boldsymbol{b(t-\tau)} - \mathbf{1})^2 - \boldsymbol{K}^2}. \tag{13}$$

The dynamical model described by Eqs. 11, 12, and 13 is summarized by the schematic in Fig. 4 in the main text.

For parameters $\boldsymbol{k}_{\mathbf{b}}$ and $\boldsymbol{k}_{\mathbf{a}}$ dictating the coarse-grained β-cat dynamics in Eq. 12, we fit to the data from single-pulse experiments directly, as shown in Supplemental Figure 6 (a) and (b). We find that $\boldsymbol{k}_{\mathbf{b}} = \mathbf{3.3} \times \mathbf{10}^{-\mathbf{3}}$ min$^{-\mathbf{1}}$ and $\boldsymbol{k}_{\mathbf{a}} = \mathbf{0.10}$ provide the best fit. Parameters for TopFlash transcription ($\boldsymbol{r}_{\mathbf{max}} = \mathbf{0.11}$ min$^{-\mathbf{1}}$, $\boldsymbol{K} = \mathbf{1.0}$, and $\boldsymbol{\tau} = \mathbf{4.0\ hrs}$) are kept the same as the model in Fig. 2. We note that the parameters $\boldsymbol{k}_{\mathbf{on}}$ and $\boldsymbol{k}_{\mathbf{off}}$ dictating the dynamics of the hidden variable have no significant impact on the fit in Supplemental Figure 6 (a) and (b) as long as $\boldsymbol{k}_{\mathbf{on}}, \boldsymbol{k}_{\mathbf{off}} \gg \boldsymbol{k}_{\mathbf{b}}$. However, their values do govern the existence and shape of the antiresonance. Supplemental Figure 6 (c) shows an example when $\boldsymbol{k}_{\text{on}} = \mathbf{8\ hr}^{-\mathbf{1}}$ and $\boldsymbol{k}_{\text{off}} = \mathbf{5\ hr}^{-\mathbf{1}}$.

We observe that the anti-resonance occurs in a parameter regime $\boldsymbol{k}_{\text{off}} < (\mathbf{1} + \boldsymbol{k}_{\mathbf{a}})\boldsymbol{k}_{\text{on}}$. This is an interesting result, as it suggests that the anti-resonance is a generic feature of the canonical Wnt pathway: it arises due to the interplay between timescales in the problem. Crucially, it depends on the ratio between in the timescales of activation and deactivation of the signaling cascade upstream of β-cat. We will now proceed to derive this bound analytically, and we show in the main manuscript that this bound matches the numerical simulations.

A periodic light input, like in the experiment, is determined by the duty cycle $\boldsymbol{\delta}$ and frequency $\boldsymbol{f}$, with period $\boldsymbol{T = 1/f}$. The duration of each light pulse is given by $\boldsymbol{\delta \cdot T}$. As such, we can write $\boldsymbol{l(t)}$ as:

$$\boldsymbol{l(t)} = \begin{cases} \mathbf{1} & \text{if } \mathbf{0} \leq \mathbf{mod}(\boldsymbol{t}, \boldsymbol{T}) < \boldsymbol{\delta T}, \\ \mathbf{0} & \text{if } \boldsymbol{\delta T} \leq \mathbf{mod}(\boldsymbol{t}, \boldsymbol{T}) < \boldsymbol{T}. \end{cases}$$

(14)

For this analysis it is convenient to consider a quasi-steady state where the system has reached a periodic solution. Let us denote the solution for $\boldsymbol{a(t)}$ in the "light on" and "light off" states as $\boldsymbol{a}_{\text{on}}(\boldsymbol{t})$ and $\boldsymbol{a}_{\text{off}}(\boldsymbol{t})$, respectively:

$$\boldsymbol{a(t)} = \begin{cases} \boldsymbol{a}_{\text{on}}(\boldsymbol{t}) & \text{if } \mathbf{0} \leq \mathbf{mod}(\boldsymbol{t}, \boldsymbol{T}) < \boldsymbol{\delta T}, \\ \boldsymbol{a}_{\text{off}}(\boldsymbol{t}) & \text{if } \boldsymbol{\delta T} \leq \mathbf{mod}(\boldsymbol{t}, \boldsymbol{T}) < \boldsymbol{T}. \end{cases}$$

(15)

Writing the initial condition as $\boldsymbol{a(0) = a_0}$, we can solve Eq. 11 to obtain:

$$\begin{aligned} \boldsymbol{a}_{\text{on}}(\boldsymbol{t}) &= \mathbf{1} - (\mathbf{1} - \boldsymbol{a_0})\, \boldsymbol{e}^{-\boldsymbol{k}_{\text{on}}\boldsymbol{t}}, \\ \boldsymbol{a}_{\text{off}}(\boldsymbol{t}) &= \boldsymbol{a}_{\text{on}}(\boldsymbol{\delta T})\, \boldsymbol{e}^{-\boldsymbol{k}_{\text{off}}(\boldsymbol{t}-\boldsymbol{\delta T})}. \end{aligned}$$

(16)

We can solve for $\boldsymbol{a_0}$ by demanding periodicity:

$$\boldsymbol{a}_{\text{on}}(\mathbf{0}) = \boldsymbol{a}_{\text{off}}(\boldsymbol{T}) = \boldsymbol{a_0},$$

(17)

which yields:

$$\boldsymbol{a_0} = \frac{\boldsymbol{e}^{\boldsymbol{k}_{\text{on}}\boldsymbol{\delta T}} - \mathbf{1}}{\boldsymbol{e}^{(\boldsymbol{k}_{\text{on}}\boldsymbol{\delta} + (\mathbf{1}-\boldsymbol{\delta})\boldsymbol{k}_{\text{off}})\boldsymbol{T}} - \mathbf{1}}.$$

(18)

First, let us consider the behavior of the model at low frequencies $\boldsymbol{T \sim 1/k_{\mathrm{b}}}$. This represents the "left side" of Supplemental Figure 6 (c). In this regime, we can ignore the comparatively fast changes in $\boldsymbol{a(t)}$ and replace it in Eq. 12 with $\boldsymbol{l(t)}$:

$$\frac{\mathbf{d}\boldsymbol{b}}{\mathbf{d}\boldsymbol{t}} \approx \boldsymbol{k}_{\mathbf{b}}\left(\mathbf{1} - \frac{\boldsymbol{b(t)}}{\mathbf{1} + \boldsymbol{k}_{\mathbf{a}}\boldsymbol{l(t)}}\right).$$

(19)

Again, we consider the quasi-steady state and denote $\boldsymbol{b(t)}$ piecewise as:

$$\boldsymbol{b(t)} = \begin{cases} \boldsymbol{b}_{\text{on}}\boldsymbol{(t)} & \text{if } \boldsymbol{0 \le \mathbf{mod}(t,T) < \delta T,} \\ \boldsymbol{b}_{\text{off}}\boldsymbol{(t)} & \text{if } \boldsymbol{\delta T \le \mathbf{mod}(t,T) < T.} \end{cases}$$

(20)

This Eq. 19 is separable, and we can solve for $\boldsymbol{b}_{\text{on}}\boldsymbol{(t)}$ and $\boldsymbol{b}_{\text{off}}\boldsymbol{(t)}$ directly by integrating. We obtain:

$$\begin{aligned} \boldsymbol{b}_{\text{on}}\boldsymbol{(t)} &= \boldsymbol{(1+k_{\mathrm{a}})\left(1 - e^{-\frac{k_{\mathrm{b}}t}{1+k_{\mathrm{a}}}}\left(1 - \frac{b_0}{1+k_{\mathrm{a}}}\right)\right)}, \\ \boldsymbol{b}_{\text{off}}\boldsymbol{(t)} &= \boldsymbol{1 + (b_{\mathbf{on}}(\delta T) - 1)e^{-k_{\mathrm{b}}(t-\delta T)}}, \end{aligned}$$

(21)

where we have denoted the initial condition $\boldsymbol{b(0) = b_0}$. We can solve for $\boldsymbol{b_0}$ by demanding that $\boldsymbol{b_0 = b_{\mathbf{off}}(0) = b_{\text{on}}(T)}$, which yields:

$$\boldsymbol{b_0 = 1 + \frac{k_{\mathrm{a}}\left(e^{\frac{k_{\mathrm{b}}\delta T}{1+k_{\mathrm{a}}}} - 1\right)}{e^{k_{\mathrm{b}}(1-\delta)T + \frac{k_{\mathrm{b}}\delta T}{1+k_{\mathrm{a}}}} - 1}}.$$

(22)

Next, we can compute the mean β-cat during one cycle:

$$\boldsymbol{\frac{1}{T}\int_0^T b(t)\mathrm{d}t = 1 + k_{\mathrm{a}}\delta - \frac{k_{\mathrm{a}}^2}{k_{\mathrm{b}}T}\left(\frac{\left(1 - e^{-k_{\mathrm{b}}(1-\delta)T}\right)\left(e^{\frac{k_{\mathrm{b}}\delta T}{1+k_{\mathrm{a}}}} - 1\right)}{e^{\frac{k_{\mathrm{b}}\delta T}{1+k_{\mathrm{a}}}} - e^{-k_{\mathrm{b}}(1-\delta)T}}\right)},$$

(23)

which is strictly decreasing as a function of frequency. This, intuitively, explains the decreasing TopFlash expression as the frequency increases in this regime. We note, however, that the functional behavior of TopFlash expression may not simply reflect the mean β-cat level during one period -- it depends on the specific TopFlash dynamics chosen. We find that a non-linear Hill-type activation (Eq. 13) strengthens the downward slope, as periods of high β-cat achieved during long pulses are "rewarded" with comparatively more TopFlash expression. Since in the low-frequency regime we have decreasing TopFlash expression as a function frequency, an "anti-resonant frequency" will appear only if TopFlash increases as we go to higher frequencies.

Next, we consider the high-frequency limit. More precisely, we consider the regime where $\boldsymbol{T \ll 1/k_{\mathrm{b}}}$. This is the "right side" of Supplemental Figure 6 (c). In this regime, the duration of each light pulse is short compared to the timescale in which significant changes occur in β-cat levels. Hence, we can solve for small oscillations of β-cat $\boldsymbol{\Delta b(t)}$ around a constant value $\widetilde{\boldsymbol{b}}$:

$$\boldsymbol{b(t) = \widetilde{b} + \Delta b(t),\ \Delta b(t) \ll \widetilde{b}.} \tag{24}$$

To first order in $\boldsymbol{\Delta b(t)}$, the β-cat dynamics in Eq. 12 can be written as:

$$\frac{\mathbf{d}\boldsymbol{\Delta b(t)}}{\mathbf{d}\boldsymbol{t}} = \boldsymbol{k}_{\mathrm{b}}\left(\mathbf{1} - \frac{\widetilde{\boldsymbol{b}}}{\mathbf{1} + \boldsymbol{k}_{\mathrm{a}}\boldsymbol{a(t)}}\right), \tag{25}$$

which is a separable equation. We can solve it directly by substituting our solution for $\boldsymbol{a(t)}$ (Eq. 16) and integrating. Denoting $\boldsymbol{\Delta b(t)}$ piecewise as:

$$\boldsymbol{\Delta b(t)} = \begin{cases} \boldsymbol{\Delta b}_{\mathrm{on}}\boldsymbol{(t)} & \text{if } \mathbf{0} \le \mathbf{mod}\boldsymbol{(t,T)} < \boldsymbol{\delta T}, \\ \boldsymbol{\Delta b}_{\mathrm{off}}\boldsymbol{(t)} & \text{if } \boldsymbol{\delta T} \le \mathbf{mod}\boldsymbol{(t,T)} < \boldsymbol{T}, \end{cases} \tag{26}$$

and imposing the initial condition $\boldsymbol{b(0)} = \widetilde{\boldsymbol{b}}$, we obtain:

$$\begin{aligned} \boldsymbol{\Delta b}_{\mathrm{on}}\boldsymbol{(t)} &= \boldsymbol{k}_{\mathrm{b}}\left[\boldsymbol{t} - \frac{\widetilde{\boldsymbol{b}}}{(\mathbf{1}+\boldsymbol{k}_{\mathrm{a}})\boldsymbol{k}_{\mathrm{on}}}\mathbf{log}\left(\frac{\boldsymbol{e}^{\boldsymbol{k}_{\mathrm{on}}\boldsymbol{t}}(\mathbf{1}+\boldsymbol{k}_{\mathrm{a}}) - (\mathbf{1}-\boldsymbol{a}_{\mathbf{0}})\boldsymbol{k}_{\mathrm{a}}}{\mathbf{1}+\boldsymbol{a}_{\mathbf{0}}\boldsymbol{k}_{\mathrm{b}}}\right)\right], \\ \boldsymbol{\Delta b}_{\mathrm{off}}\boldsymbol{(t)} &= \boldsymbol{\Delta b}_{\mathrm{on}}\boldsymbol{(\delta T)} + \boldsymbol{k}_{\mathrm{b}}\boldsymbol{(t-\delta T)} - \boldsymbol{k}_{\mathrm{b}}\left[\frac{\widetilde{\boldsymbol{b}}}{\boldsymbol{k}_{\mathrm{off}}}\mathbf{log}\left(\frac{\boldsymbol{e}^{\boldsymbol{k}_{\mathrm{off}}\boldsymbol{(t-\delta T)}} + \boldsymbol{k}_{\mathrm{a}}\boldsymbol{a}_{\mathrm{on}}\boldsymbol{(\delta T)}}{\mathbf{1}+\boldsymbol{k}_{\mathrm{a}}\boldsymbol{a}_{\mathrm{on}}\boldsymbol{(\delta t)}}\right)\right]. \end{aligned} \tag{27}$$

Like before, we can solve for $\widetilde{\boldsymbol{b}}$ by demanding periodicity $\boldsymbol{\Delta b}_{\mathrm{on}}\boldsymbol{(0)} = \boldsymbol{\Delta b}_{\mathrm{off}}\boldsymbol{(T)} = \widetilde{\boldsymbol{b}}$. This yields:

$$\begin{aligned} \widetilde{\boldsymbol{b}} = (\mathbf{1}+\boldsymbol{k}_{\mathrm{a}})\,\boldsymbol{k}_{\mathrm{off}}\,\boldsymbol{k}_{\mathrm{on}}\,\boldsymbol{T}\Bigg[&(\mathbf{1}+\boldsymbol{k}_{\mathrm{a}})\,\boldsymbol{k}_{\mathrm{on}}\,\mathbf{log}\left(\frac{\boldsymbol{e}^{\boldsymbol{k}_{\mathrm{off}}\,\boldsymbol{(1-\delta)T}} + \boldsymbol{a}_{\mathrm{on}}\boldsymbol{(\delta T)}\boldsymbol{k}_{\mathrm{a}}}{\mathbf{1}+\boldsymbol{a}_{\mathrm{on}}\boldsymbol{(\delta T)}\boldsymbol{k}_{\mathrm{a}}}\right) \\ &+\boldsymbol{k}_{\mathrm{off}}\,\mathbf{log}\left(\frac{\boldsymbol{e}^{\boldsymbol{k}_{\mathrm{on}}\,\boldsymbol{\delta T}}\,(\mathbf{1}+\boldsymbol{k}_{\mathrm{a}}) - (\mathbf{1}-\boldsymbol{a}_{\mathbf{0}})\boldsymbol{k}_{\mathrm{a}}}{\mathbf{1}+\boldsymbol{a}_{\mathbf{0}}\boldsymbol{k}_{\mathrm{a}}}\right)\Bigg]^{-\mathbf{1}}. \end{aligned} \tag{28}$$

Next, we consider the functional behavior of $\widetilde{\boldsymbol{b}}$ with frequency. For anti-resonance to exist, we know that $\widetilde{\boldsymbol{b}}$ must increase as the frequency increases towards the high-frequency limit. Formally, one needs to show that (i) $\widetilde{\boldsymbol{b}}$ is monotonic $\boldsymbol{T}$ and (ii) determine the sign of $\mathbf{d}\widetilde{\boldsymbol{b}}/\mathbf{d}\boldsymbol{T}$. If $\mathbf{d}\widetilde{\boldsymbol{b}}/\mathbf{d}\boldsymbol{T} < \mathbf{0}$, anti-resonance occurs. Here, we take a shortcut by considering the limits of $\boldsymbol{T} \to \mathbf{0}$ and $\boldsymbol{T} \to \infty$. These are:

$$\begin{aligned} \lim_{T\to 0} \tilde{b} &= \frac{k_{\text{off}}(1-\delta) + (1+k_{\text{a}})k_{\text{on}}\delta}{k_{\text{off}}(1-\delta) + k_{\text{on}}\delta} \\ \lim_{T\to\infty} \tilde{b} &= \frac{1+k_{\text{a}}}{1+k_{\text{a}}(1-\delta)} \end{aligned} \tag{29}$$

For anti-resonance, we require that:

$$\lim_{T\to 0} \tilde{b} > \lim_{T\to\infty} \tilde{b}. \tag{30}$$

Substituting the limits in Eq. 29 into the above inequality, we obtain the condition:

$$\boxed{k_{\text{off}} < (1+k_{\text{a}})k_{\text{on}},} \tag{31}$$

in agreement with results from the numerical simulations.

**Supplementary Table 1:** T-test results from TopFlash (TF) and β-catenin (β-cat) traces from Fig 1D

| Names | P-value TF | P-value β-cat | P-value TF Stars | P-value β-cat Stars |
|---|---|---|---|---|
| 6HR and 9HR | 8.65E-20 | 5.94E-05 | *** | *** |
| 6HR and 15HR | 2.19E-61 | 4.76E-12 | *** | *** |
| 6HR and 18HR | 4.56E-94 | 6.43E-19 | *** | *** |
| 6HR and 21HR | 1.11E-85 | 2.46E-12 | *** | *** |
| 6HR and 24HR | 9.14E-05 | 2.60E-12 | *** | *** |
| 9HR and 15HR | 9.05E-24 | 1.06E-05 | *** | *** |
| 9HR and 18HR | 6.39E-49 | 3.20E-14 | *** | *** |
| 9HR and 21HR | 2.88E-54 | 2.41E-08 | *** | *** |
| 9HR and 24HR | 7.52E-40 | 4.73E-06 | *** | *** |
| 15HR and 18HR | 1.93E-08 | 2.92E-06 | *** | *** |
| 15HR and 21HR | 6.50E-23 | 0.00795 | *** | ** |
| 15HR and 24HR | 1.12E-95 | 0.90173 | *** | n.s. |
| 18HR and 21HR | 4.16E-08 | 0.05985 | *** | n.s. |
| 18HR and 24HR | 1.86E-139 | 3.08E-07 | *** | *** |
| 21HR and 24HR | 2.06E-124 | 0.00334 | *** | ** |

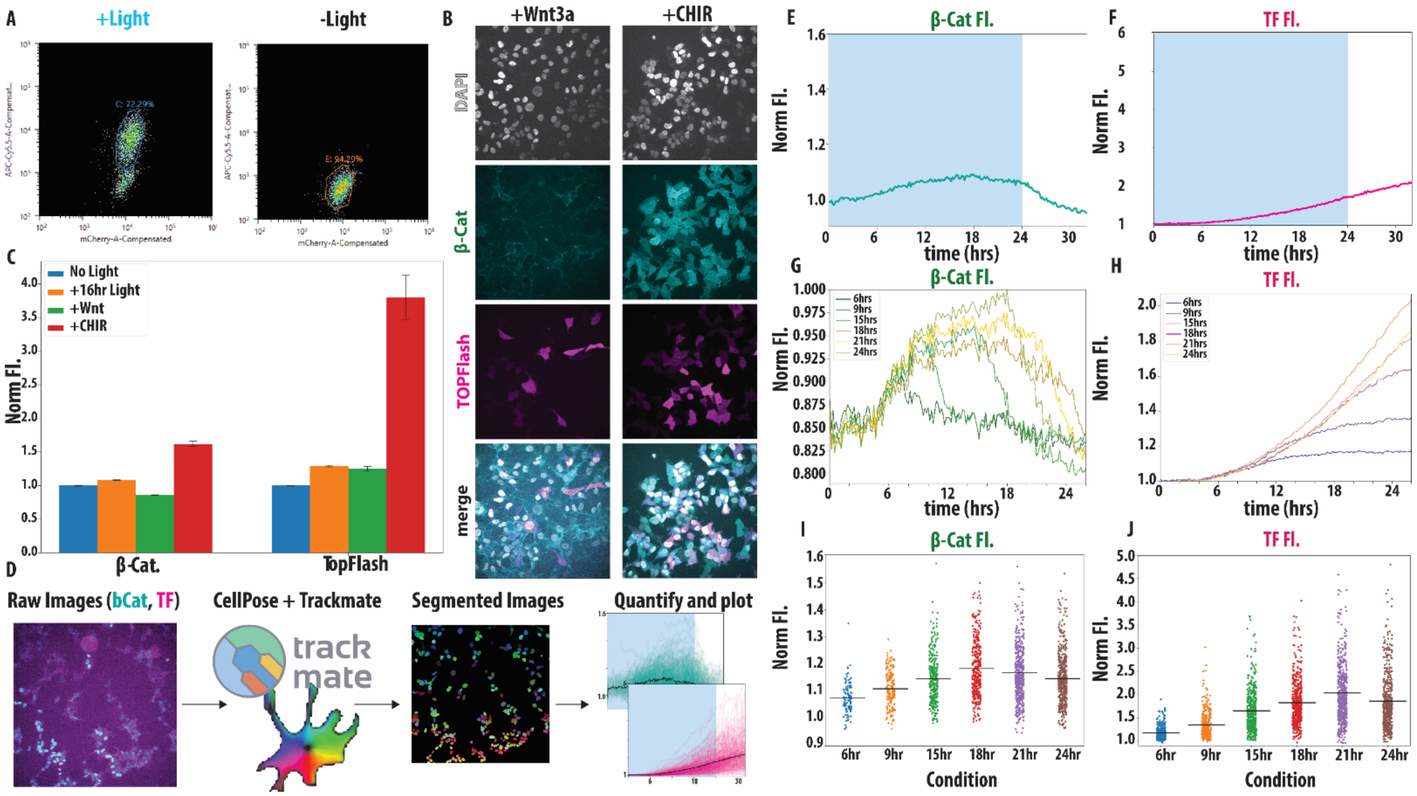


**Supplementary Figure 1:** (A) FACS data for Wnt I/O cell line with (+)/without (-) 24hrs of light exposure. Axes are of β-catenin (mCherry) v. TopFlash (APC-Cy5.5) (B) Live cell imaging of CRISPR tdmRuby3-β-cat, lentiviral 8X-TOPFlash-tdIRFP and DAPI, 16 hours after adding CHIR99021 (+CHIR) and Wnt3a (+Wnt3a). (C) Quantifications of β-catenin and TopFlash fluorescence for no light, +16hrs light, +Wnt3a and +CHIR, where the mean of each condition was normalized to the mean of the no light condition and error bars represent SEM. (D) Flow for cell segmentation and heatmap generation from experimental data, using CellPose add TrackMate. From left to right: magenta (β-catenin) and cyan (TopFlash) images are passed into CellPose + Trackmate for segmentation and tracking. Example image of CellPose segmentation is shown under “segmented images”, where the different colors correspond to the cell’s segmentation ID. Final images under “quantify and plot” shows quantification of β-catenin and TopFlash. (E) Mean fluorescent intensity (MFI) of β-catenin in the 24hrs light on condition, normalized to light off β-catenin. (F) MFI of TopFlash in the 24hrs light on condition, normalized to light off TopFlash. (G) Population mean MFI of β-catenin from live, single cell traces in indicated conditions, normalized to light off β-catenin. Initial drop in fluorescence is due to media bleaching. (H) Population mean MFI of TopFlash from live, single cell traces from indicated conditions, normalized to light off TopFlash. (I) Jitterplot of β-catenin mean nuclear fluorescent intensity (MFI) at the maximum intensity point in continuous light exposure conditions. Each point represents a single cell and the black line represents the mean of the population. (J) Jitterplot of TopFlash mean nuclear fluorescent intensity (MFI). Each point represents a single cell and the black line represents the mean of the population.

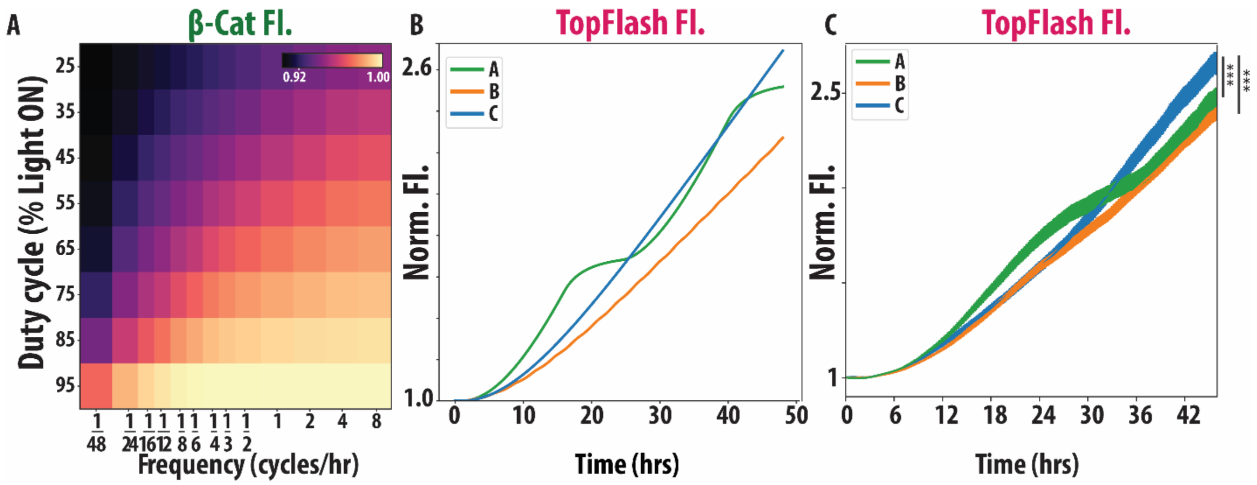


**Supplementary Figure 2:** (A) ODE model generated heatmap of endpoint β-catenin MFI for various combinations of duty cycle and frequency conditions. (B) ODE model generated TopFlash MFI dynamic traces for points A, B and C from Figure 2F. (C) Experimentally generated TopFlash MFI dynamic traces for point A, B and C from Figure 2F. Error bars represent standard error of the mean (SEM).

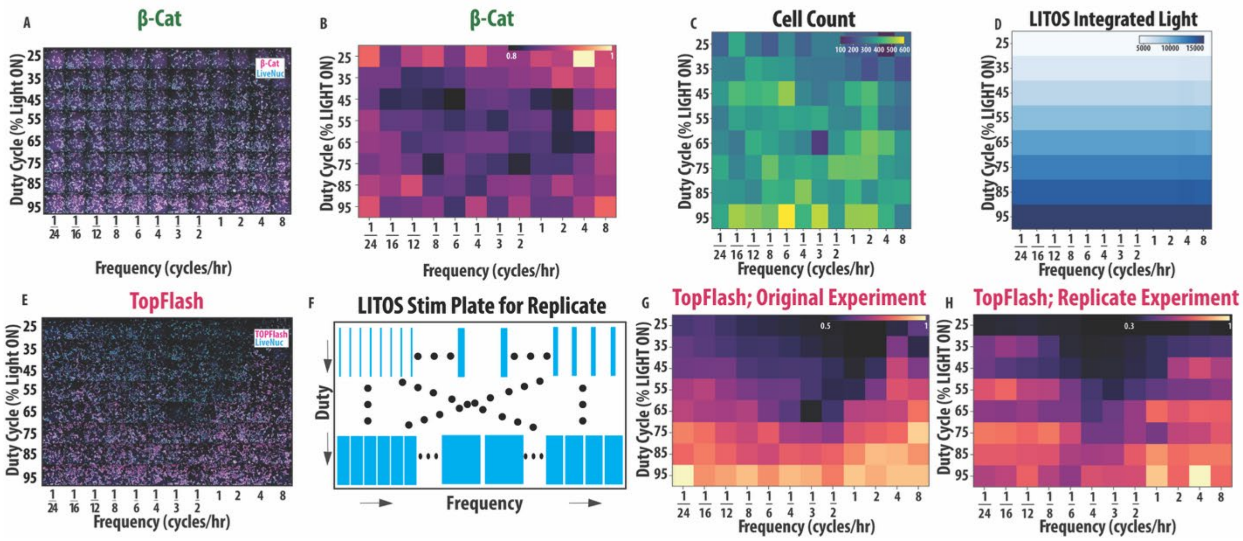


**Supplementary Figure 3:** (A) β-catenin and LiveNuc stained Wnt I/O HEK293T cells imaged post duty cycle and frequency experiment, ran for 48 hours. (B) Heatmap of end point β-catenin mean fluorescent intensity (MFI) post duty cycle and frequency experiment where values were normalized to the maximum β-catenin MFI. (C) Heatmap of total cell count for each duty cycle and frequency condition. (D) Heatmap of the total integrated amount of light for each duty cycle and frequency condition, obtained from the LITOS conditions used in the experiment. (E) TopFlash and LiveNuc stained Wnt I/O HEK293T cells imaged post duty cycle and frequency experiment, ran for 48 hours. (F) Rearranged LITOS stimulation plate set up for the replicate duty cycle and frequency experiment. (G) Heatmap of end point TopFlash MFI from our original duty cycle and frequency experiment, normalized to the maximum TopFlash MFI. (H) Heatmap of end point TopFlash MFI from our replicate duty cycle and frequency experiment, normalized to the maximum TopFlash MFI.

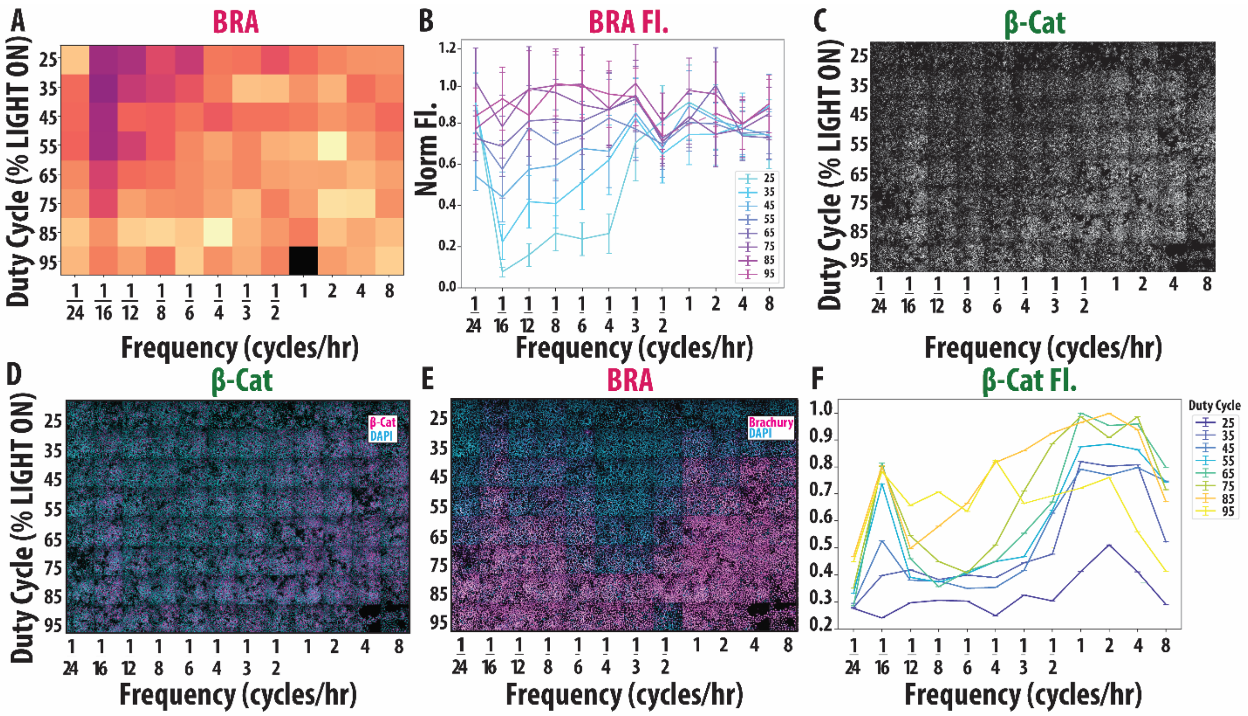


**Supplementary Figure 4:** (A) Heatmap of endpoint BRA in H9 Wnt I/O cells following duty cycle and frequency stimulation, ran for 48 hours. This experiment replicates the conditions shown in Fig. 5 but extends the stimulation duration from 24 hours to 48 hours. (B) Line plot of endpoint BRA in H9 Wnt I/O cells following duty cycle and frequency stimulation, ran for 48 hours. This experiment replicates the conditions shown in Fig. 5 but extends the stimulation duration from 24 hours to 48 hours. Error bars represent standard error of the mean (SEM). (C) Qualitative images of the end point β-catenin fluorescence post LITOS illumination (N = 862-3176 cells, 6 biological replicates per condition). (D) Qualitative images of the end point β-catenin and DAPI fluorescent stain post LITOS illumination. (E) Qualitative images of the end point BRA and DAPI fluorescent stain post LITOS illumination. (F) Error bar plot of end point β-catenin MFI. Error bars represent SEM.

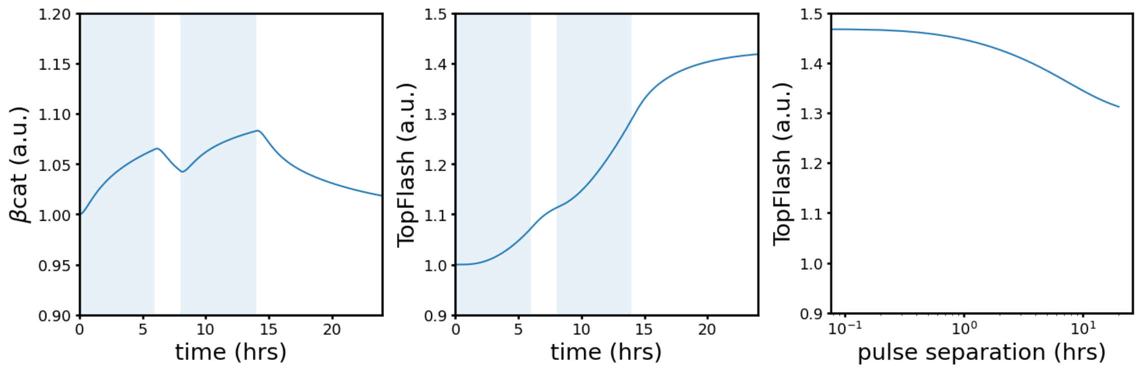


(a) (b) (c)

**Supplemental Figure 5:** (a) and (b) show the β-cat and TopFlash traces for two 6-hr pulses with a pause duration of 2 hrs. Figure (c) shows the final TopFlash expression level (at $\boldsymbol{t} = \mathbf{48}$ hrs) after two 6-hr pulses with varying pause duration. Blue shading indicates opto-Wnt activation during these time intervals.

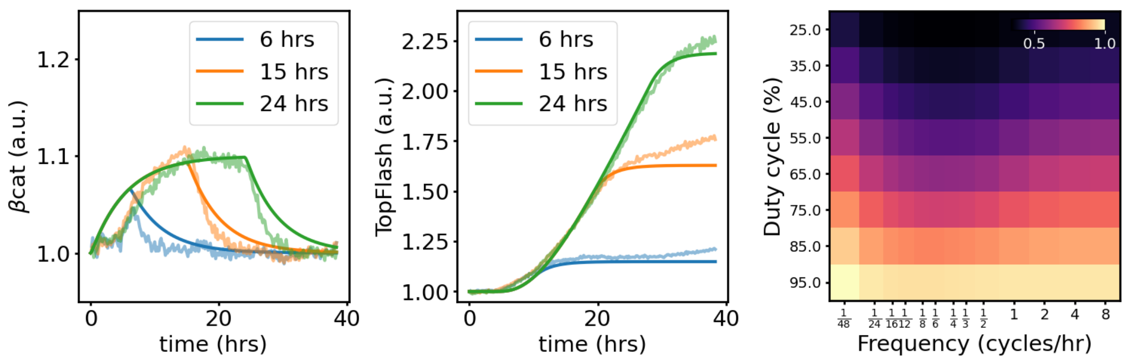


(a) (b) (c)

**Supplemental Figure 6:** Figures (a) and (b) show traces from the hidden-variable model of β-cat and TopFlash for three single-pulse conditions. Dynamics in this regime are dictated by $\boldsymbol{k}_{\mathbf{b}}$ and $\boldsymbol{k}_{\mathbf{a}}$. Low-alpha lines show experimental data. Figure (c) shows an example of the anti-resonance when $\boldsymbol{k}_{\text{on}} = \mathbf{8\ hr^{-1}}$ and $\boldsymbol{k}_{\text{off}} = \mathbf{5\ hr^{-1}}$.

**Supplementary Movie 1:** Video of clonal HEK293T Wnt I/O under 24hrs of continuous light exposure, delivered as 100ms long pulses every 2 minutes followed by an 8hr relaxation period. *Top Left:* Wnt I/O HEK293T cell line β-catenin fluorescent channel, imaged every 10 minutes. In the absence of 405nm blue light stimulation, β-catenin fails to accumulate in the nucleus due to no optogenetic activation of opto-Wnt tool. *Top Right:* Wnt I/O HEK293T cell line TopFlash fluorescent channel, imaged every 10 minutes. In the absence of 405nm blue light stimulation, TopFlash fails to accumulate due to no optogenetic activation of opto-Wnt tool. *Bottom Left:* Wnt I/O HEK293T cell line β-catenin fluorescent channel, imaged every 10 minutes. In the presence of 405nm blue light stimulation, β-catenin accumulates in the nucleus due to optogenetic activation of opto-Wnt tool. *Bottom Right:* Wnt I/O HEK293T cell line TopFlash fluorescent channel, imaged every 10 minutes. In the presence of 405nm blue light stimulation, TopFlash accumulates due to optogenetic activation of opto-Wnt tool.

**SI References**


1. Lee E, Salic A, Krüger R, Heinrich R, Kirschner MW (2003) The roles of APC and Axin derived from experimental and theoretical analysis of the Wnt pathway. *PLoS Biol* 1:e10. https://doi.org/10.1371/journal.pbio.0000010.
2. Tan CW, Gardiner BS, Hirokawa Y, Layton MJ, Smith DW, Burgess AW (2012) Wnt signalling pathway parameters for mammalian cells. *PLoS ONE* 7:e31882. https://doi.org/10.1371/journal.pone.0031882.
3. de Man SM, Zwanenburg G, van der Wal T, Hink MA, van Amerongen R (2021) Quantitative live-cell imaging and computational modeling shed new light on endogenous WNT/CTNNB1 signaling dynamics. *eLife* 10:e66440. https://doi.org/10.7554/eLife.66440.
4. Harris TJC, Peifer M (2005) Decisions, decisions: beta-catenin chooses between adhesion and transcription. *Trends Cell Biol* 15:234–237. https://doi.org/10.1016/j.tcb.2005.03.002.
5. Goentoro L, Kirschner MW (2009) Evidence that fold-change, and not absolute level, of beta-catenin dictates Wnt signaling. *Mol Cell* 36:872–884. https://doi.org/10.1016/j.molcel.2009.11.017.
6. Bauer M, Graf IR, Ngampruetikorn V, Stephens GJ, Frey E (2017) Exploiting ecology in drug pulse sequences in favour of population reduction. *PLoS Comput Biol* 13:e1005747. https://doi.org/10.1371/journal.pcbi.1005747.
